# SAXS DATA BASED GLYCOSYLATED MODELS OF HUMAN ALPHA-1-ACID GLYCORPROTEIN, A KEY PLAYER IN HEALTH, DISEASE AND DRUG CIRCULATION

**DOI:** 10.1101/2024.02.01.578488

**Authors:** Nidhi Kalidas, Nagesh Peddada, Kalpana Pandey, Ashish

**Author notes:** Address correspondence to: Ashish, PhD, CSIR-Institute of Microbial Technology, Sec 39A Chandigarh INDIA 160036. Nidhi Kalidas: Department of Biology, University of Marburg, Marburg 35043 Germany. Nagesh Peddada, PhD: The Center for the Genetics of Host Defense, UT Southwestern MedicalCenter, Dallas-TX 75235 USA.

## Abstract

Plasma Alpha-1-glycoprotein (AGP) binds diverse drugs, its isoforms and their levels vary significantly in acute phases of health. Relative binding of drugs to AGP and albumin has been used to model their release profiles, structural insights on glycosylated form of AGP will certainly improve estimations. Main challenges are: 1)heavy glycosylation of AGP (~50% of its mass), 2) isoforms of primary structure co-exist, and 3) extreme heterogeneity in the glycan motifs have been reported. Our small angle X-ray scattering (SAXS) data on plasma extracted AGP showed interparticulate effect from 283-313 K which disappeared irreversibly upon further heating to 343K. Using ALPHAFOLD2 server, the protein only portion could be modelled but its theoretical SAXS profile did not match acquired experimental data. Using mass spectra-based information, we attached glycan motifs at different known sites to compute four models of fully glycosylated AGP. Importantly, calculated SAXS profiles of our glycosylated models agreed well with the experimental data. Docking runs revealed that *in silico* interaction of different drugs when varied using unglycosylated *vs*. glycosylated model as receptor. Finally, we propose that our SAXS based models of glycoprotein are better representation of the molecule and should be considered for structure-based analysis and/or estimations.

## Introduction

Levels of human Alpha-1-glycoprotein (AGP), its chemical polydispersity, and its differential interactions with varied versions with drugs have been reviewed, concluding that this protein is a critical indicator of health and regulator of drug circulation in body ^1,2^. It is also known as Orosomucoid (ORM) and belongs to Lipocalin family of proteins ^2^. AGP is a heavily glycosylated protein, with about 50% of its molecular mass from attached glycans ^3,4^. Theoretical mass of unglycosylated AGP is about 23.6 KDa, and native human (glycosylated) AGP migrates close to 45 KDa in SDS-PAGE ^5^. Analysis showed that negatively charged sialic acid groups dominate the branched glycosylation which makes its pI to be 2.7-3.2 ^5,6^. Publications showed that serum levels of AGP increase several folds during acute phase response as systemic response to inflammation ^7^. Detailed investigation of the glycosylation pattern brought forth that they change in relation to the induced inflammation and these changes regulate function in platelets and neutrophils ^4–9^. Expression of human AGP is mediated by three adjacent genes: while AGP-A encodes the F1, F2 and S variants, AGP-B and AGP-B’ encode A variant ^10,11^. The precursor AGP protein is a 201 amino acid protein with a secretary N-terminal signal peptide of 18 residues which gets cleaved during processing to result in final version of protein with 183 amino acids ^12^. Distribution of these variants in different populations, their ability to dampen different immune conditions/responses have been reported and highlight polydisperse constitution of AGP in plasma ^13,14^. It is documented that the molar ratios of Σ(F1, F2 and S) forms to A form is about 3:1 to 2:1 in blood of healthy individuals ^15^, and this ratio gets skewed in acute phase conditions like breast cancer and chronic inflammatory disease ^16,17^. Analytical mass spectrometry (MS) data analysis of plasma from lymphoma, melanoma and ovarian cancer patients reported that the molar ratio of Σ(F1, F2 and S): A forms were about 8:1 ^18^. These sets of reports conclude that not only total AGP levels varied under disease conditions, but also the relative fraction of the isoforms changed. Based on the primary structure, signal sequence processed AGP has five N-linked glycan positions: Asn15, Asn38, Asn54, Asn75 and Asn85 ^19^. Interestingly, F1 and F2 forms have 38^th^ position as Gln, and S form has Arg in the same position ^20^. A form of AGP has about 20 substitutions compared to other forms and these mutations in the primary structures also result in decrease in the number of attached glycans. Furthermore, attached glycans contain different degrees of branching (bi- to tetra-antennary) ^6^. Complicating further, terminating sugars are known to have high degrees of diversity in their constitution from sialic acid to fucose, and their incorporation varies in people having inflammation ^21,22^.

Thus, along with variation in the levels of isoforms at primary structure level, the number of glycans, their branching and identity of terminating sugar increases the overall heterogeneity of AGP molecules in our body,as confirmed from MS experiments ^23–27^. Several efforts have been made to decipher the glycosylation pattern in AGP ^23,24,27,28^. A quick look at the ElectroSpray Ionization mass spectrometric Time of Flight (ESI-qTOF-MS) profiles published by Baerenfaenger and Meyer of human plasma AGP brings out the extremely high level of polydispersity in the chemical constitution of this protein in real life ^28^. Analysis of data provided several peaks in the range of 35 to 38.5 KDa for the intact mass of AGP. Additionally, the authors processed MS data from disialyted AGP protein samples to comprehend their observations from fully glycosylated AGP. Using pooled plasma samples, these authors found 90, 101 and 64 different glycan constitutions for the F1, S and A versions, respectively ^28^. Comparison of AGP constitution from individual donors showed significant variations in fucosylation and branching of glycans. Similar analysis of AGP glycoprofiling revealed that levels and constitution varies under diabetic or hyperglycemic conditions during critical illness ^24^. In 1992, Treuheit and co-workers published putative molecular masses of glycans at five sites in human plasma AGP and their possible branching using Fast Atom Bombardment (FAB) and Matrix Assisted Laser Desorption (MALDI) TOF mass spectrometric data ^27^. For glycosylation sites1 through 5 in AGP, 3, 3, 5, 11 and 6 different molecular masses could be interpreted, reconfirming extensive variation in the branching patterns at individual sites. Deglycosylation attempts resulted in some deconvolution of the data, but the primary conclusion from these experiments was that there is a high degree of variability in glycan structures. Importantly, the latter publication provided some vital clues about molecular masses of the glycan attachments at the five sites of AGP ^27^. These numbers aided us in our current study to search for representative motifs in building working models of this biomedically relevant plasma protein ^2,5,7,9,23^.

Over last few decades, binding of different drug or drug-like candidates to plasma AGP has been of interest as this glycoprotein plays critical role in drug uptake, distribution, deposition and action ^1^. Binding of several drugs with diverse chemical structures to AGP which correlated their clinical implications, have been published and reviewed ^1,29–32^. Interestingly, training on quantitative biochemical data on free vs. bound drug in plasma, attempts have been made to predict binding and release profiles of drugs to AGP ^31,33–36^. Drug binding profiles to A and F/S variants have been studied ^37^. Based on observed binding profile of 35 diverse drugs, and using crystal structures of unglycosylated AGP, a 3D quantitative structure-activity relationship (QSAR) model was developed ^37,38^. Additionally, relative binding of drugs to plasma albumin and AGP was used to predict drug profiles which upon comparison with experimental data indicated that for better predictions, variations in protein and its glycosylation levels and identity must be considered for converging results with experiments ^2,9,36,39^. Today, with the availability of reliable insight into the structure of protein receptor, range of force-fields, and increased prowess of modern computing systems, one can easily embark on reliable computer aided drug design to its release profile in our bodies.

One important factor missing from achieving realistic structure-based prediction of binding profiles of small drugs to AGP is a working model of a glycosylated version of this protein which is based on some reliable experimental data on the glycan decorated version of AGP. To date, crystal structures of A and F1/S variants are available −/+ different drug like compounds. Crystal structures of AGP A variant in complex with disopyramide and amitriptyline revealed aromatic interactions occur between drug and sidechains of Phe112 and Phe49 (PDB IDs 3APW and 3APV) ^40^. Interestingly, in the co-crystal structures of unglycosylated AGP, different bound drug molecules were seen to be bound in the same pocket of the protein without significant changes in rest of the structure. Another non-specific molecule,chlorpromazine known to experimentally bind native AGP with a differential mode, was also seen in the same cavity as others (PDB ID 3APX) ^40^. Additionally, modelling studies and available co-crystals were used to understand various ways in which these molecules could bind unglycosylated AGP and apply corrections to fit experimentally observed results ^41–44^. Experimentally, binding of drugs to AGP forms were done using biophysical techniques like Circular Dichroism (CD), fluorescence, photoaffinity labelling ^44–46^. For all experimental work, while polydisperse population of natural glycosylated AGP, theoretical predictions were done using unglycosylated models. To provide researchers with a reliable model of glycosylated AGP, we acquired and analyzed SAXS datasets from human plasma derived AGP. Here, we added previously suggested sugar moieties on a structure of unglycosylated AGP to generate models of this glycoprotein. Importantly, theoretical SAXS profiles of our glycosylated models agree better with the experimental SAXS data. Coordinates of this glycosylated model was used to dock some drugs /organic moieties known to bind AGP. Results presented in this work imply that many drugs show differential binding behavior to unglycosylated vs. glycosylated model of AGP.

## Material and Methods

### SAXS Data Acquisition and Processing

Native human AGP extracted pooled human plasma was purchased from ABCAM [CatLog No. AB109930]. Commercially supplied lyophilized protein sample was dissolved in PBS pH 7.4. The solution was filtered through 0.2-micron filter, and protein concentration in the filtrant was estimated to be ~3.2-3.5 mg/ml based on band intensity of single band close to 45 KDa from sample vs. that for standard protein (BSA) in SDS-PAGE.This sample and matched buffer were used to acquire SAXS profiles from solution and buffer. All SAXS experiments were done at SAXSpace Instrument in CSIR-IMTECH equipped with sealed tube source in line collimation (Anton Paar Austria) ^47–49^. Scattered X-rays were recorded on 1D Mythen detector (Dectris). Samples and buffer were exposed toX-rays for one hour each. Datasets were collected at temperature 283 to 343 K at intervals of 10K, each for one hour. Then, experimental temperature was decreased from 343 to 283 K in steps of 10K and data was collected for one hour each. Programs used for data collection to processing are listed in **Supplementary Table T1**. Direct desmearing was done using the beam profile and SAXSQuant program which provided SAXS dataset profiles representing pinhole camera profile ^48^. Then, buffer subtraction was done from the solution SAXS profiles at different temperatures in auto mode. Finally, we obtained intensity profiles, I(s) as a function of momentum transfer vector, s where s = 4π(sin θ/λ) with units in 1/nm. SAXS profiles of the solute i.e., AGP at different temperatures were analyzed using ATSAS suite of programs v 3.0.1 and 3.0.2 ^50^ and ATSAS online portal. Shape parameters like Radius of Gyration (R_g_), maximum linear dimension (D_max_) and molecular weight (MOW) were estimated using the SAXS profiles ^51,52^. CRYSOL program was used to compute theoretical SAXS profiles from the protein (and glycoprotein) models and compare them with the experimental SAXS datasets ^53^. The SAXS datasets for AGP at 283 and 343 K, (and solved 3D models of glycosylated AGP) are available at SASBDB database under submission IDs SASDPG4 and SASDPH4, respectively.

### Modelling of Glycosylated Models

First using primary structure of human AGP (UniProt P02763), its unglycosylated model was generated using SWISS-MODEL ^54^ and AlphaFold2 server ^55^. Residues1-183 of the post-signal sequence of AGP was considered for structure building. Options were selected to use monomer or multimer status during prediction of AGP model(s). Using AlphaFold2 server, five similar models of monomeric unglycosylated AGP were predicted based on Lipocalin fold. Best scored model was used to generate model for the glycoprotein by using it for input template at GlyProt server ^56^. This server enables attachment of manually selected N-glycan motif to the 3D structure of protein at specified N-glycan site in non-clashing conformation. Glycan motifs used and their positions are mentioned later in the results section. Four combinations of glycan moieties were placed to generate four models of glycoprotein AGP. Computed SAXS profiles from the glycoprotein models were compared with the experimental datasets. Normal mode analysis of glycosylated models was done using ELNEMO server ^57^. For the latter calculations, default parameters were employed to compute five low frequency modes.

### Cavity Analysis for Docking Studies

Using best-scoring models of unglycosylated and glycosylated AGP, pockets or cavities in 3D structures were calculated. Two web based servers were used: FPOCKET ^58^ (https://bioserv.rpbs.univ-paris-diderot.fr/services/fpocket/) and GHECOM ^59^ (https://www.google.com/search?client=firefox-b-d&q=GHECOM+). Both servers considered PDB format models as input, and had option to use scalable size of probe to explore pockets in the 3D structure being analyzed. While FPOCKET runs on Voronoi tessellation algorithm, GHECOM is a grid-based search which computes and uses mathematical morphology of the target structure.

### Docking of Small Molecules on Models of AGP

Different small molecules known to bind AGP (and some selected randomly) were docked on proteinogenic and glycoprotein models of AGP using MTiAutoDock v 1.0 (https://mobyle.rpbs.univ-paris-diderot.fr/cgi-bin/portal.py#forms::MTiAutoDock) ^60^. Input files were one ligand at a time in mol2 format, and PDB structure of target. Docking considered the complete structure of the target protein and flexible conformations of ligand with rotation across allowed bonds. Top ten poses based on low energy on target were selected for visual comparison between unglycosylated and glycosylated models.

## Results and Discussion

### SAXS Data on Native Human AGP

SAXS data was collected from human AGP at a concentration of about 3.2-3.5 mg/ml at different temperatures from 283 to 343 K and matched buffer profile was subtracted (**Figure 1** and **Supplementary Figure S1**). SAXS profiles of the AGP sample as it was heated from 283 to 343 K and then temperature was decreased back to 283 K are shown in **Figure 1A**. No upward trend in the intensity values as s approaches 0 supported complete lack of protein aggregation in the temperature range studied (**Figure 1A** and **Supplementary Figure S1**). By dividing the SAXS data profile at 343 K by that collected for 283 K, we obtained ratios of intensities at different s values (**Figure 1B**). The flatness of the trend or slope of zero to a ratio close to 1 should have reflected no change in the SAXS profiles being compared, but the Left panel indicated that there were changes in the shape of AGP molecules in vectors higher than 2 nm^−1^ (or lower than 3.14 nm) as the sample was heated to 343 K. The small peak profile at s ~ 0.8 nm^−1^ (or 7.85 nm) indicated loss of interparticulate effect upon heating since ratio of intensities remained close to 1 as s was close to 0 nm^−1^. In contrast, the ratio of profiles upon cooling the sample showed a slight dip at 0.8 nm^−1^ and unchanged slope from 2 nm^−1^ onwards implying slight changes in large dimensions and no appreciable change in smaller dimensions, as the same sample was cooled. Peak profiles of the Kratky plots of the SAXS datasets support that molecules adopt globular shapes in solution (**Figure 1C**). In correlation to comparison shown in **Figure 1B**, Guinier approximation for globular shapes showed that slope of linear fits becomes less negative at lower s values (**Supplementary Figure S2**). Corresponding, normalized Kratky plots also peaked slightly lower than sR_g_ value of 1.73. Furthermore,automatic estimation of the Pair Distribution profiles indicated negative probability for vectors beyond 6-6.5 nm for all datasets except at 323 and 333K. These three observations supported interparticulate effect or anomalous behavior in estimating longer interatomic vectors possibly originating from glycans to glycans decorating the outershell of this heavily glycosylated protein. The latter is likely explanation since sugar moieties would have density and solvation properties different from proteinogenic constitution alone. Please see the panels in **Supplementary Figure S2** and parameters listed in **Table 1** which indicate that at temperature 333 K, the anomalous behavior disappeared, and Pair Distribution could be calculated correctly in automated manner. Reversal of temperature from 343 to 283 K showed that datasets at each 10 K, remained almost identical to that at 343 K, thus only forward heating related SAXS datasets were analyzed. This concluded that unorthodox behavior seen for AGP molecules at lower temperatures is not restored by merely cooling the sample. Or, in other words, heating removes the interparticulate behavior seen at lower temperatures. Molecular mass (MOW) of the AGP molecules estimated from their SAXS data showed expected increase with temperature (**Table 1**). At 323 and 333 K, analysis estimated a molecular mass of ~25 to 45 kDa, whose average about the mass estimated for human AGP for MS ^28^. Moreover, the molecular mass estimated for AGP based on datasets at 333 and 343 K is about 47± 3 kDa which interestingly correlates with the glycoprotein’s reported migration pattern of ~45 kDa on SDS-PAGE ^5^. The SAXS datasets for AGP at 283 and 343 K, exhibiting and lacking anomalous profile are available at SASBDB database as IDs SASDPG4 andSASDPH4, respectively.

**Figure 1.**
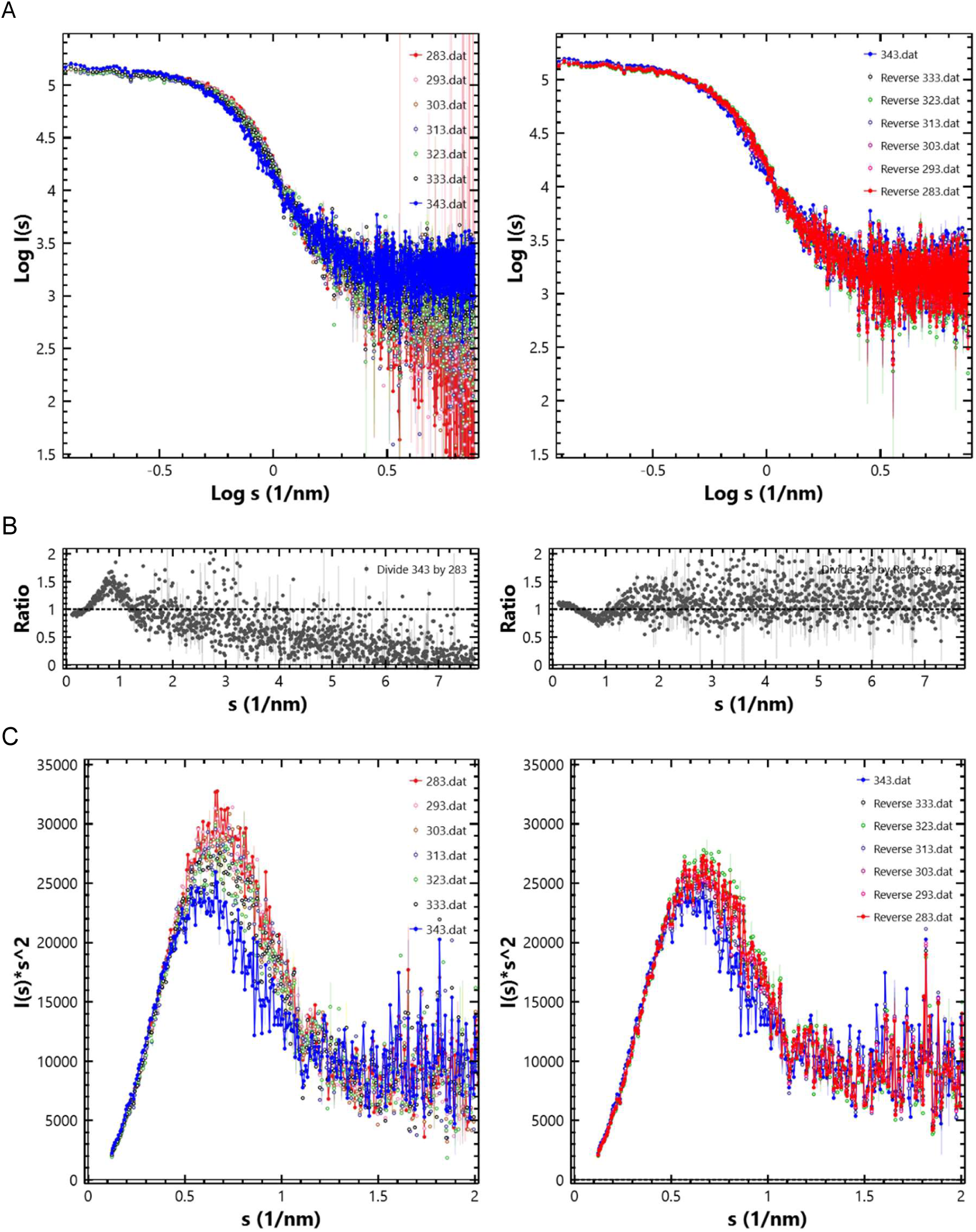
SAXS profiles from sample of AGP. (A) Solution SAXS profiles from AGP at different temperatures are shown in double Log mode, as the temperature was increased from 283 to 343 K (Left) and as the temperature was decreased from 343 to 283 K (Right). Experimental temperatures of the datasetare mentioned as legends. (B) Comparison of the datasets at 343 K with 283 K by dividing I(s) values of former by latter profile, as temperature was increased (Left) and decreased (Right). In both plots, the horizontal dotted line at relative intensity value of 1 is the expected profile if both unscaled profiles were identical. (C) Kratky plots of the SAXS datasets presented in panel A are shown here. *Additional plots of the datasets are in* ***Supplementary Figure S2***. All figures are made using ATSAS 3.0.2 data analysis software.

**Table 1:** Structural parameters deduced analysis of the SAXS datasets from sample of AGP at different temperatures are tabulated below. The Pair Distribution profiles-based parameters which significantly disagreed with the respective Guinier analysis due to presence of interparticulate effect are shown with underlines. Datasets are shown in **Figure 1**. Please, also see **Supplementary Figure S2** for analysis details.

| Temperature (K) | Guinier Analysis | Bayesian Inference | Pair Distribution |  |
| --- | --- | --- | --- | --- |
| | $R_g$ (nm) [s$R_g$ limits] | MOW (kDa) | $D_{max}$ (nm) | $R_g$ (nm) |
| 283 | 2.06 [0.25-1.24] | 28.9 | <u>8.62</u> | <u>1.82</u> |
| 293 | 2.10 [0.26-1.29] | 31.65 | <u>8.42</u> | <u>1.61</u> |
| 303 | 2.11 [0.26-1.26] | 28.9 | <u>8.06</u> | <u>1.96</u> |
| 313 | 2.16 [0.27-1.27] | 26.87 | <u>7.41</u> | <u>2.23</u> |
| 323 | 2.18 [0.27-1.27] | 24.95 | 6.22 | 2.38 |
| 333 | 2.33 [0.29-1.30] | 44.72 | 6.42 | 2.49 |
| 343 | 2.53 [0.32-1.13] | 46.65 | <u>9.03</u> | <u>2.58</u> |

### Models of AGP

There are couple of crystal structures available for the proteinogenic part of AGP, with some bound to drug candidates ^40,61^. Crystal structures of A variant in unliganded and bound to some drugs are shown in **Supplementary Figure S3**. It must be highlighted here that the unglycosylated protein in PDB submissions 3APU, 3APV and 3APW were refined as dimer in their asymmetric unit. On the other hand, structure was solved as monomer in PDB submissions 3APX and 7OUB, raising curiosity about monomer/ dimer status of glycan-less version of AGP in solution. The retention of the Lipocalin fold in the structures and little large-scale changes in the protein structures bound to ligands lend support to our hypothesis that the proteinogenic portion folds intrinsically, and post-translational modifications follow the core protein’s folding process on its surface. Additionally, the active site seen in co-crystal structures can absorb/accommodate a variety of ligands without requiring large scale readjustments in AGP. Same logic was employed by us to model this glycoprotein *i.e.,* first the proteinogenic core was modelled, then the glycan moieties were attached to the surface of protein structure. Finally, the computed SAXS from models of glycoprotein were compared with the experimental SAXS profiles to assess results. Foremost, the primary structure of human AGPwas considered for template-based modelling using FASTA format sequence as mentioned in the methods section. Theoretical mass of the protein portion of human AGP is 21.6 kDa only, which was much lower than the mass of intact glycoprotein of about 35-38 kDa ^27^. First, SWISS-MODELserver was considered, where HHblits protocol suggested 3APX as best template for AGP ^54^. Sequence identity was 89.07% and the template was for AGP2 solved with a ligand 3-(2-chloro-10H-phenothiazin-10-yl)-N, N-dimethylpropan-1-amine with a resolution of 2.2 Å ^40^. Second best template was 3KQ0 which was solved at 1.8 Å for AGP1 protein and sequence identity of 100% ^61^. This latter structure was solved bound to (2R)-2,3-dihydroxypropyl acetate. ProMod3 protocol of SWISS-MODEL server modelled 1-178 residues using template PDB ID 3APX but did not model the C-terminal “EEGES”.

ALPHAFOLD2 server was also employed to model protein part of AGP ^55^. Multiple sequence alignment with available templates and predicted local distance difference test results of the search are shown in **Supplementary Figure S4A** and **S4B**. There was lower sequence identity and thus lower certainty in the N- and C-terminal part of the models predicted by ALPHAFOLD2. Five comparable models covering all 183 residues were computed as best possible options with variations in the N- and C-terminal ends (**Supplementary Figure S4C**). The top ranked model of protein portion (of the five similar solutions) is shown in **Figure 2A**. Its theoretical SAXS was compared with the experimental SAXS data from AGP at 283 and 343 K (**Figure 2B**). (*Additionally, calculated SAXS profiles of all five solutions from ALPHAFOLD2 are compared with experimental datasets in the* ***Supplementary Figure S3D***). The theoretical R_g_ value of the five models of protein only portion was 1.7 nm or 17 Å, which was much smaller than experimentally estimated R_g_ value about 2.5-2.7 nm for AGP (**Table 1**). More importantly, the calculated SAXS profile of the protein only model did not compare well with the profiles of experimental datasets upholding that protein only model lacks similarity with molecular shape of glycosylated AGP.

**Figure 2.**
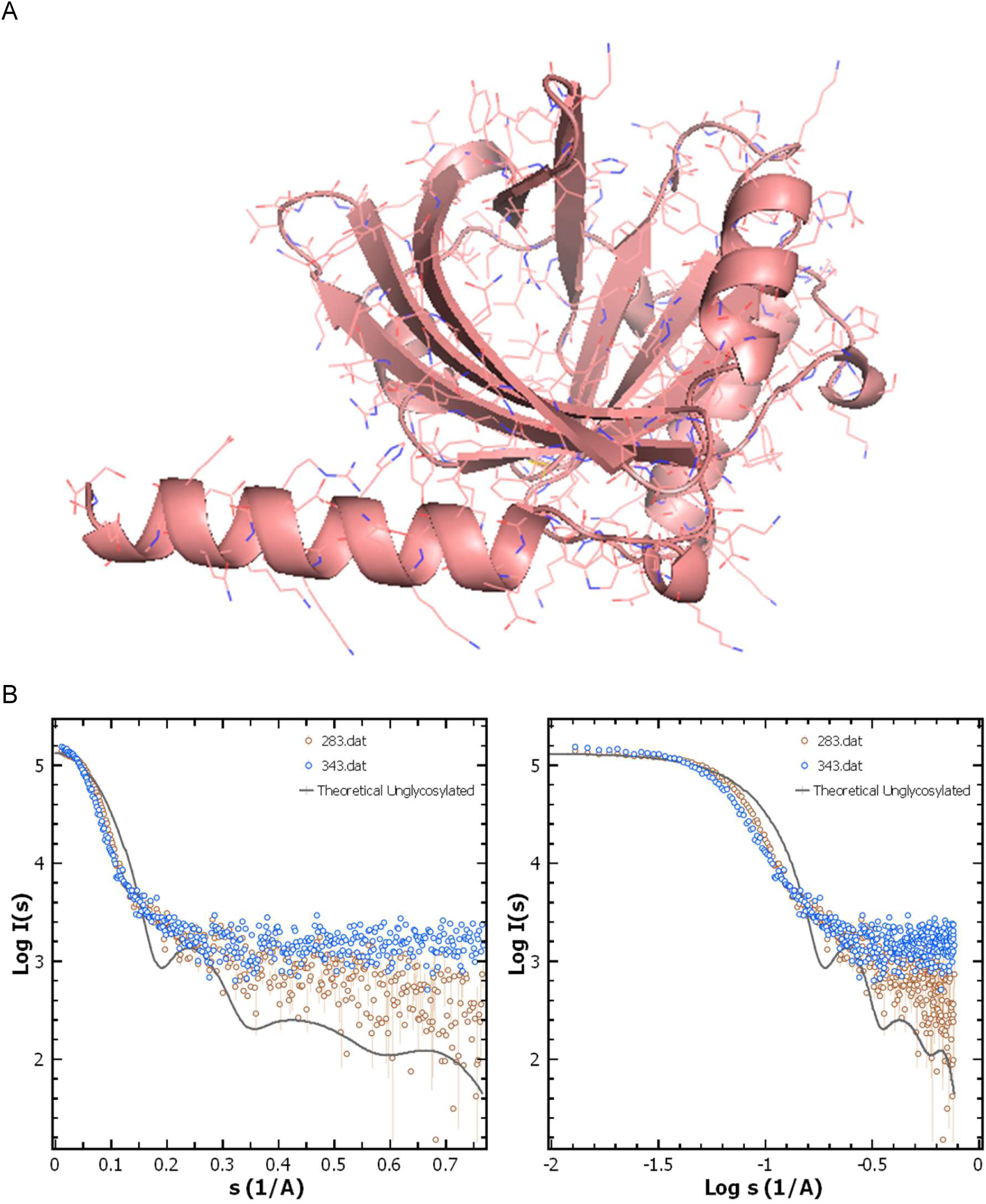
Predicted structure of protein only part of AGP. (A) Best scored model of AGP predicted from ALPHAFOLD2 server is shown here in cartoon mode. Sidechains of amino acids are shown as lines. The image was made using the PyMoL program. (B) SAXS profile computed for this model (shown as solid black line) is compared with SAXS datasets collected for 283 and 343 K. Right and Left panels are plots in Log vs. Linear mode, and double Log mode, respectively. Images of plots were made using ATSAS 3.0.1 data analysis software.

There are five Asn residues in AGP which are sites of glycan attachments during post-translational processing: 15^th^, 38^th^, 54^th^, 75^th^ and 85^th^ residues (Treuheit et al., 1992). Importantly, all these five Asn residues are on the surface of the model of the proteinogenic portion. In the model shown in **Figure 2A**, side-chain torsion angles of Asn15 are 60°, 340°, 160° and 260°; Asn38 are 220°, 340°,160° and 260°; Asn54 are 300°, 20°, 160° and 260°; Asn75 are 300°, 20°, 140° and 260°; and Asn85 are 300°, 140°, 160° and 260°. The χ^1^- χ^4^ torsion angles indicate that the sidechains are extended away from protein structure, and thus can be attached with glycan moieties without steric clashes. In 1992, Treuheit *et al*. reported detailed analysis of the five glycosylation sites in human AGP ^27^. They analyzed MS profiles of different desialylated glycoforms of protein and digested purified glycopeptides to get estimation of possible glycan structure(s) and their mass(es) at different sites in AGP. For site 1 to 5, MS data suggested 3, 3, 5, 11 and 6 possible glycan moieties with average masses about 3948 ± 263, 2855 ± 366, 3826 ± 340, 4957 ± 266 and 3487 ± 372Da, respectively. (*These average mass values of glycans are estimated by presuming their equal propensity*). Additionally, these authors interpreted the glycans to be complex structured: site 1 - bi- to tri-antennary, site 2 – bi- to tetra-antennary, site 3 - bi- to tetra-antennary, site 4 – tri- to tetra-antennary, and site 5 - bi- to tetra-antennary, plus differential sugar groups in them. This indicated the high level of chemical polydispersity in the glycans present in AGP and it complicated generation of models of glycoprotein. Upon searching GLYPROT database, we found glycan motifs not precisely same, but with masses closer to the estimated average values (site 1 – glycan IDs 8959 and 8533, both with mass 4029 Da; site 2 – glycan IDs 9119, 9085, 9073, all with mass 2862 Da, site 3 – glycan ID 8951, mass 4108 Da; site 4 – glycan ID 9232, mass 5274 Da; and site 5 – glycan ID 8996, mass 3664 Da). Their different structures and branching can be seen in **Supplementary Figure S5**. Thus, four combinations were used for the five sites: glycan motif IDs were 8959/8533, 9119/9085/9073, 8951, 9232, and 8996. Due to this, compared to the hypothesized average mass per site, the error in mass value of the motif used was: site 1 – 80 Da, site 2 – 7 Da, site 3 – 281 Da, site 4 – 316 Da, and site 5 – 176 Da. Except for the site 4, these errors in selecting glycan motifs per site were less than estimated variation between masses measured at each site experimentally by MS data ^27^. The total mass of the glycans to be connected was about 19.9 kDa which along with the proteinogenic portion of 21.6 kDa suggested the molecular mass of glycoprotein AGP close to 41 kDa. By considering the four above mentioned combinations of glycan motifs, four models of AGP were generated by attaching specified glycan motifs on 3D model of protein (**Figure 3A**). Theoretical SAXS profiles of the four models of glycoprotein were compared with the experimental SAXS datasets at 283 and 343 K (**Figure 3B**). Compared to the SAXS profile computed for protein only model, the computed profiles of glycoprotein models better fitted experimental data at both SAXS and WAXS regions. This upheld that the glycosylated models are better representation of the actual structure of native AGP in solution.

**Figure 3.**
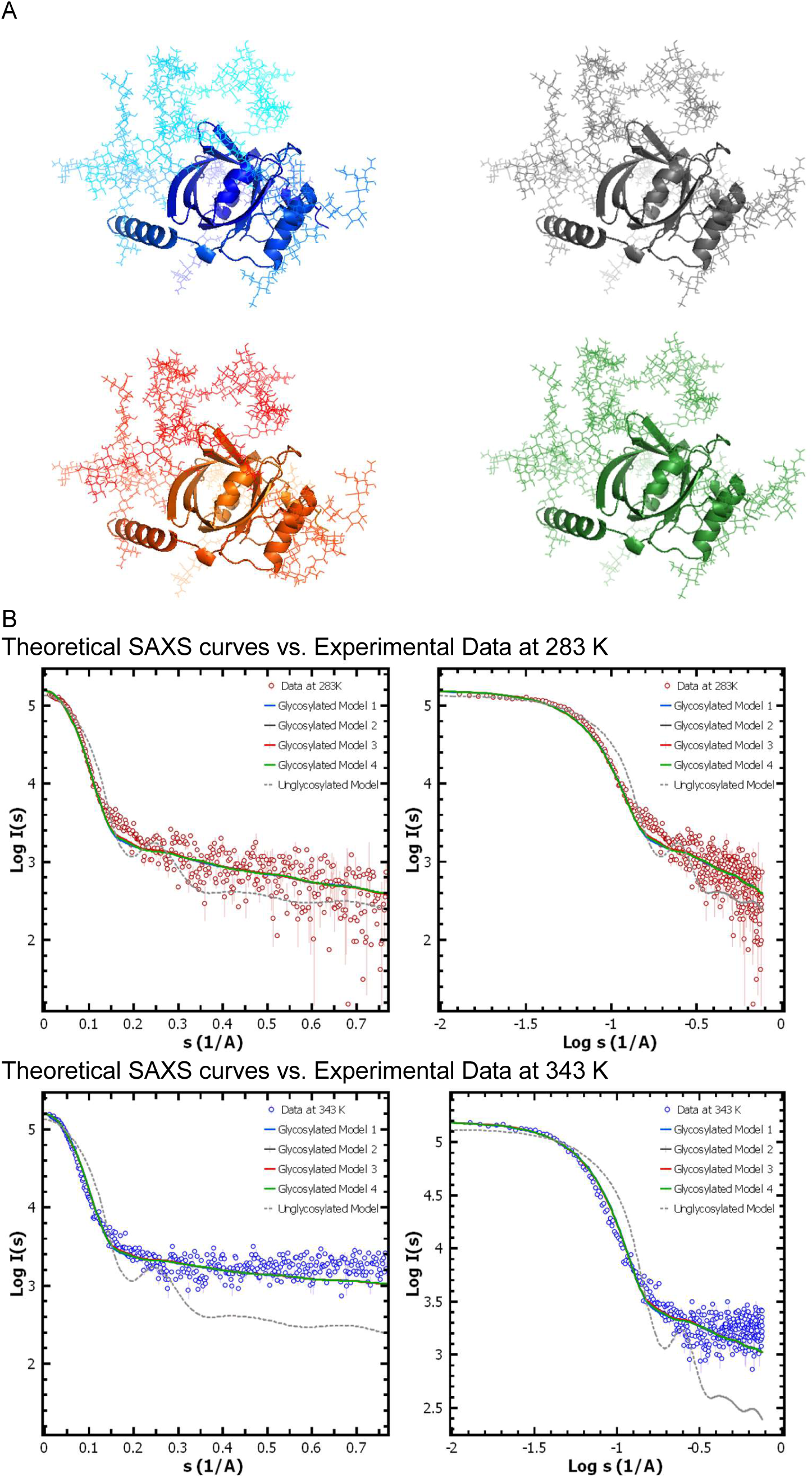
Glycosylated models of AGP. (A) Four models of AGP with glycan moieties (lines) attached to protein (cartoon) as described in script are shown here. Images were made using the PyMol program. (B) Computed SAXS profiles of the four models (colored solid lines as models in panel A) are compared with SAXS datasets at 283 and 343 K. Right and Left panels are plots in Log vs. Linear mode, and double Log mode, respectively. Black dotted line shows the computed SAXS profile for protein only model. Images of plots were made using ATSAS 3.0.1 data analysis software.

### Understanding the solution shape of glycosylated model of AGP

To perceive extent of inherent motion and altered hydrodynamic profile due to attachment of glycans, normal mode analysis was employed to compute low frequency motions in the glycosylated models of AGP (**Figure 4A**). The ligated glycans in the models also vibrated about the proteinogenic core which helped in perceiving how the hydrodynamic behavior of the glycoprotein AGP are significantly dominated by the glycan moieties. Theoretical envelope R_g_ was increased by 53% from 17.1 to 26 Å or 1.7 to 2.6 nm for protein to glycoprotein, respectively. For the protein only and glycosylated model, the envelope surface area and volume increased from 4133 and 7729 Å^2^, and 2.83E+04 and 7.69E+04 Å^3^, respectively. This implied that addition of glycan moieties, in agreement with previous MS and our SAXS data, led to increase in surface area and volume by almost 87 and 172%, respectively. The branched glycan moieties attached to Asn sidechains appeared to wobble in conical volumes around the protein core. This meant that interacting molecules or drugs which are known to bind AGP would primarily see glycan moieties rather than proteinogenic portion ^1,2,40^, Based on known co-crystals of unglycosylated AGP with small molecules, some theoretical protocols have been made to predict/assess binding profiles to this protein ^41–44^. Yet, experimental binding using different biophysical and biochemical methods were done mostly using glycosylated version of the AGP ^44–46^. Using the model of glycosylated model of AGP, we wanted to explore differences in binding profiles of drug like molecules vs. using only proteinogenic core.

**Figure 4.**
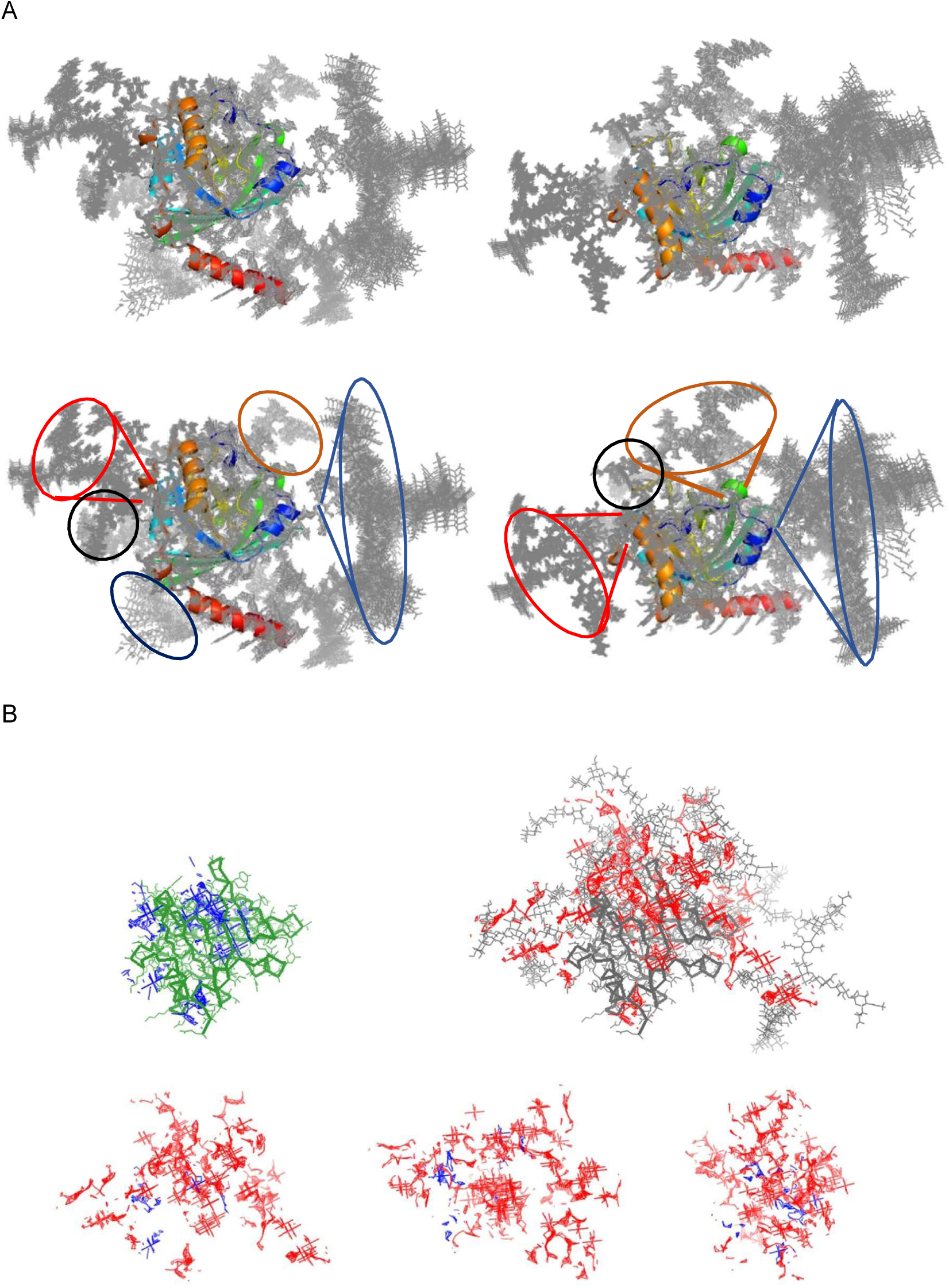
Computed low frequency motion in glycoprotein models and cavities in two models of AGP. (A) Normal mode calculations-based results are shown here. Five low frequency motions of all four glycoprotein models are shown here. Protein is shown in cartoon mode and coloring is blue to red from N- to C-terminal, and attached glycan moieties are shown as grey lines. Circles drawn in the lower panels (same view as above) emphasize how glycans may move around protein and significantly increase the hydrodynamic properties of AGP. (B) Results of pocket location in the structural models using FPOCKET server and a probe radius of 6 Å are shown here. Blue and red lines represent the pockets in the protein only (green) and glycoprotein (grey) models, respectively. The lower panel shows rotated views of the two sets of pockets superimposed in spatial reference to AGP models. The images of models and pockets were generated using the PyMoL program.

Using FPOCKET and GHECOM servers, and models of unglycosylated and glycosylated version of AGP, we calculated probable druggable pockets in the 3D structures ^58,59^. Results of pocket location in the structural models using FPOCKET server and a probe radius of 6 Å, and their correlation with primary sequence and respective surface area are shown in **Figure 4B** and **Supplementary Figure S6**, respectively. While in the protein only model, there were four or five main plausible pockets including the site most seen in the co-crystals of AGP, there were several additional putative binding pockets in the glycosylated model. In the case of latter, the pockets were in protein part, as well as moieties composed of both protein and glycans, and in glycan moieties suggesting wide variation of binding pockets are available in glycoprotein version. Superimposition of the placement of pockets in 3D space of two models helped in visualizing how additional pockets theoretically emerge in glycosylated model around the protein core. Importantly, similar results were obtained by using GHECOM server and probe radius of 6 Å (**Supplementary Figure S7**). To explore the binding possibility of slightly bigger drug molecules like alkaloids *etc*., the probe radius was increased to 10 Å which showed expected trend of many additional pockets in the glycan decorated model of AGP. These results upheld a thought of using glycosylated model for drug docking studies and compare profiles with protein only docking target.

### Docking profiles of small molecules on models of AGP

Using model of glycosylated AGP which agreed better with the experimental SAXS data, we performed *in silico* docking of few drugs known to bind AGP and few other drug moieties to see if they bind differentially than to protein only model (**Figure 5**). These were: known drugs - Aripiprazole, Staurosporine, Progesterone, Propranolol, Disopyramide, Verapamil, Warfarin and moieties – Tuberine, Ostreogrycin A, Doxorubicin, Sabarubicin and Sucrose. While understanding mechanism of binding of Aripiprazole to AGP was attempted via extrinsic Cotton effects, the magnitudes of the induced circular dichroism did not correlate concluding mechanism of binding of Aripiprazole to this glycoprotein remained unexplained (Nishi et al., 2019). Staurosporine, and its hydroxylated forms UCN-01 and UCN-02, are kinase inhibitors that bind to both AGP1 and AGP2, and though co-crystal of UCN-01 with AGP2 is known, solution NMR suggests ligand binding induced conformational changes which are yet to be delineated for these antitumor compounds ^62^. Crystal structure of A variant of AGP bound to UCN01 (PDB ID 7OUB) showed an RMSD of only 0.45 Å over 150 C^α^ residues with another structure of AGP bound to polyglycol (PDB ID 3APU). Clearly, results from NMR experiments did not agree with the crystal structures. A lower affinity of chlorpromazine and progesterone for the molten globule state of AGP (which is reported to form at pH 4.5 and 40°C) suggested that ligand(s) may be released near the negative surfaces of biological membranes ^63^. Additionally, it has been proposed that interaction of AGP with reverse micelles induced a unique β-sheet to α-helices which reduced binding capacity for the drugs, chlorpromazine and the steroid hormone, progesterone to AGP ^64^. Interestingly, AGP is known for exhibiting chiral preference while binding to propranolol with bound concentration for (S)-propranolol than (R)-propranolol ^65^. Similarly, AGP as stationary phase showed preferential enantiomeric binding to verapamil vs. its main metabolite norverapamil ^66^. Furthermore, ultrafast affinity extraction and peak profiling were used with AGP as matrix to study binding profiles for several model drugs (i.e., chlorpromazine, disopyramide, imipramine, lidocaine, propranolol, and verapamil) ^67^. These experiments showed modest association constants to AGP in the range of 10^4^-10^6^M^−1^ at pH 7.4 and 37°C. One study reported that the free disopyramide in serum was affected by alteration in levels of AGP in congestive heart failure and acute myocardial infarction ^68^. Another study showed binding of chlorpromazine, imipramine, propranolol, and warfarin increased by 13-76% to AGP from systemic lupus erythematosus cases, while disopyramide decreased by 21-25%, in compared to normal cases ^69^. Binding profiles of anticoagulant warfarin to AGP and haptoglobin have been correlated to some extent with the heterogeneity in the glycan components of proteins which showed that increased antennae branching and terminal fucosylation, reduced drug-binding affinity ^70^. Thus, we selected these drugs for comparative docking profiles on the models of AGP, protein only and glycosylated. Additionally, we randomly selected some drugs/moieties for comparative docking.

**Figure 5.**
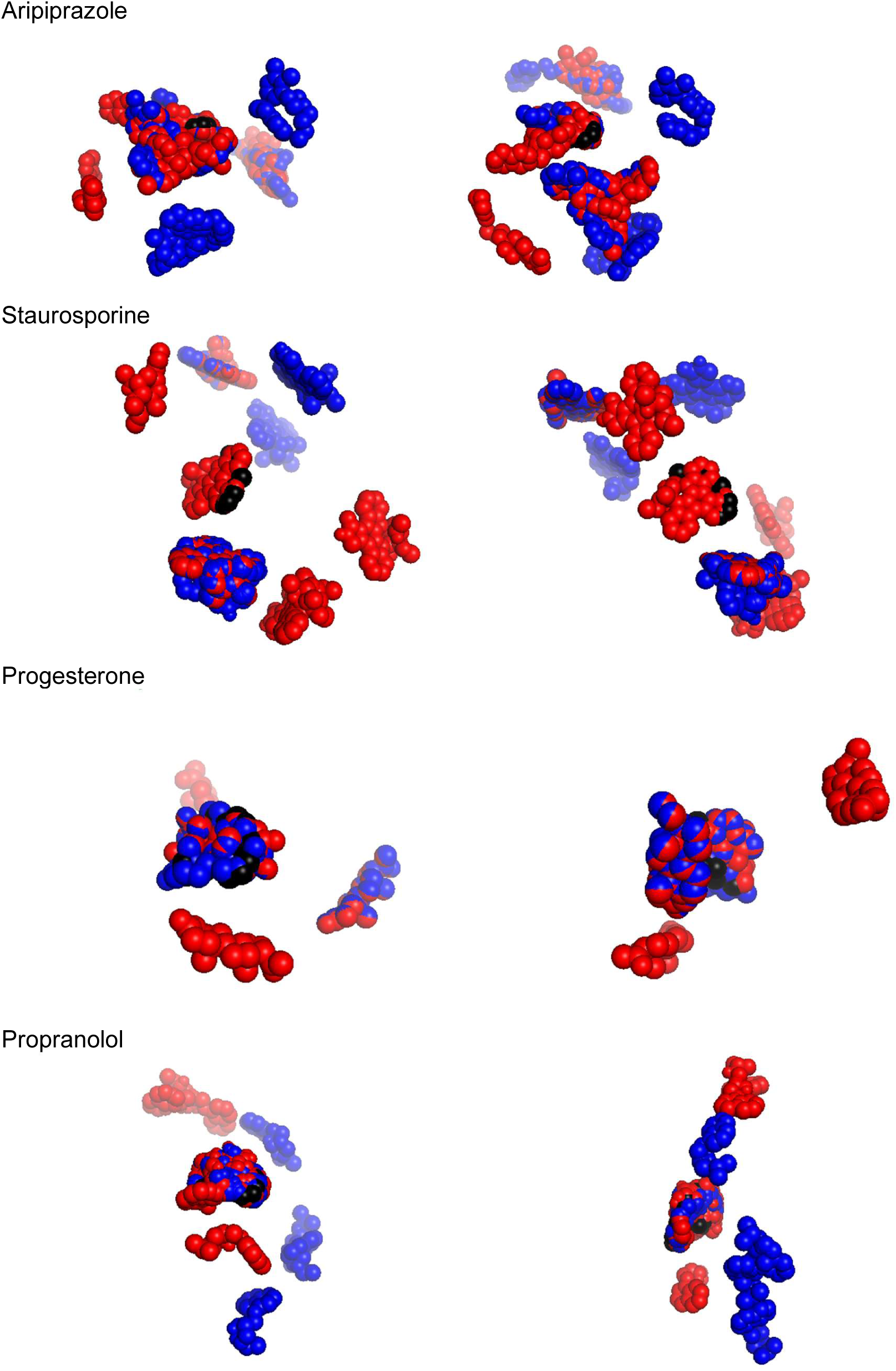
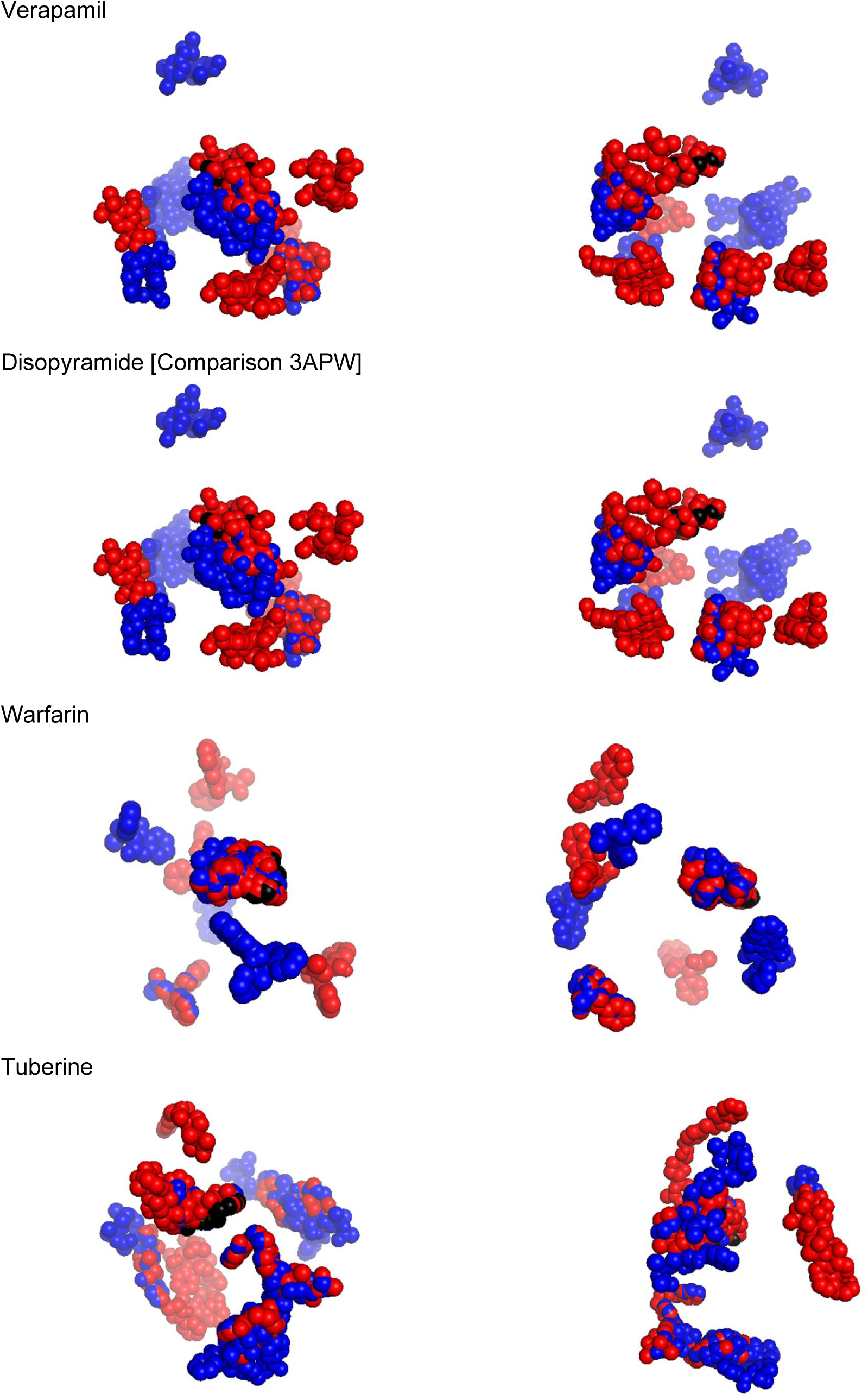
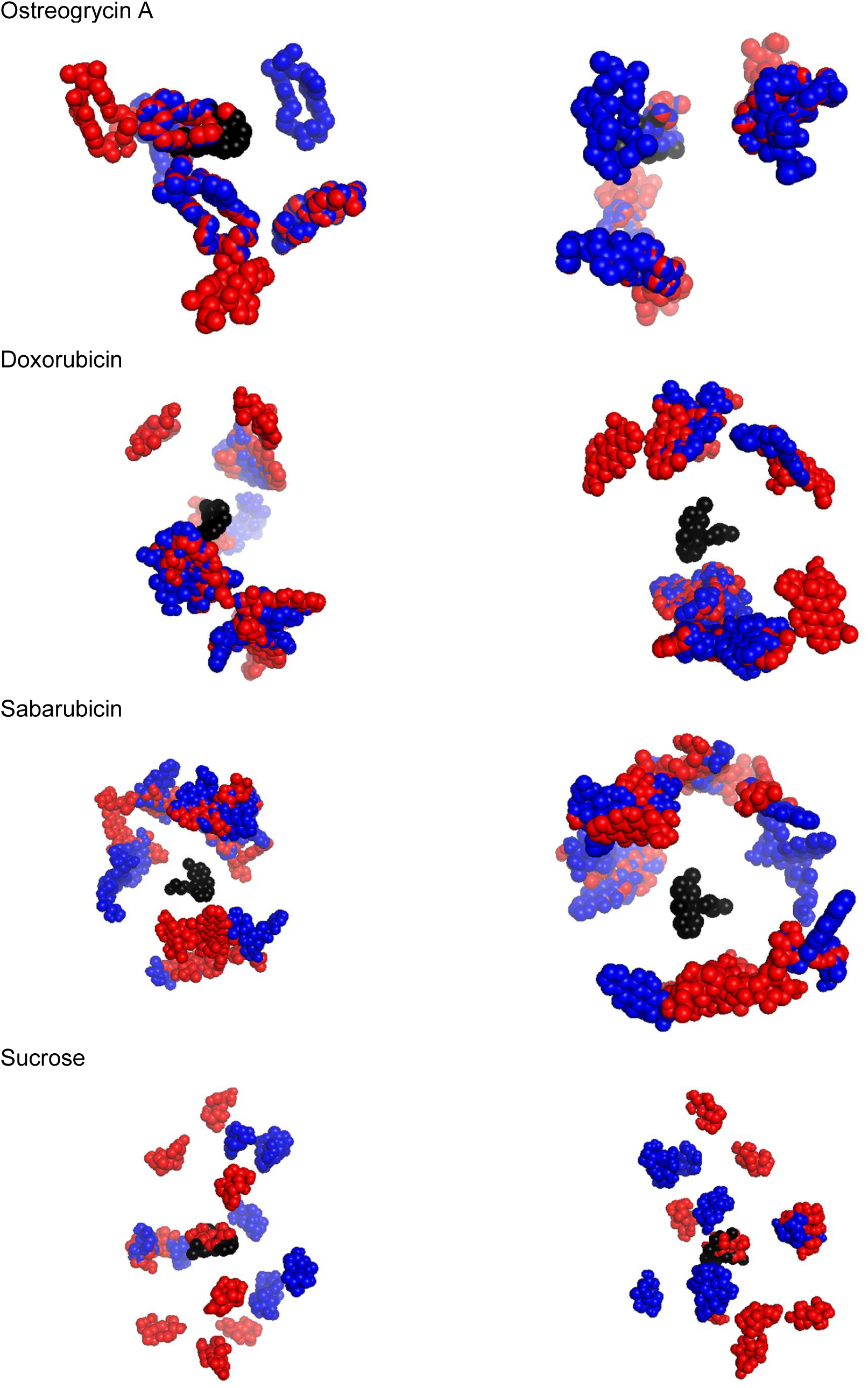
Comparison of the results from docking drugs on the models of AGP. Drug or organic moiety docked models of AGP are mentioned above the panels. Two columns show two rotated views of the top ten docked poses of drugs on protein model (blue cpk) and glycoprotein model (red cpk). The position and pose of disopyramide as seen in the co-crystal structure PDB ID 3APW is shown as black cpk. The glycoprotein (and the protein) structures in the same rotation are shown in the Supplementary Figure S8. The images were generated using the PyMoL program.

Considering flexible or rotatable structures of drugs / moieties and static 3D models of protein only and glycosylated AGP, docking experiments were done. It is pertinent to mention here that there are some publications which argue multisite binding profiles of drugs or small molecules to single site as the concentration range is narrowed ^71–73^. This was a valid consideration for AGP, since differing ligands were always resolved bound in the same cavity in various co-crystals, and little detectable change in rest of the structure of AGP. Of course, all crystal structures were set-up with unglycosylated AGP. Thus, to explore multi-site binding theory for AGP ^72^ and as mentioned in methods section, for each docking run, top ten dock poses on protein structures were considered for analysis. In **Figure 5**, ligand poses docked on protein model are shown in blue cpk, and those docked on the model of glycosylated model of AGP are shown in red cpk. The protein part in both models of AGP are superimposed which allowed comparison of dockposes of ligand(s) and shown in **Supplementary Figure S9**. The glycosylation moieties are shown as grey lines, protein part is shown as grey cartoon, and last 6 residues at C-terminal are shown in orange. In Figure 5, the two columns show how the ligands interacted with the surfaces of the protein models. No drug or moiety screened with glycan only portion, but mostly interacted with protein only or surfaces jointly formed by protein and glycan. Disopyramide in its position as bound to AGP in crystal structure is shown as black cpk to compare how far our docked poses are in both models. For almost all ligands, some blue and red docked positions were similar, even the pose of the ligand was comparable, but for all cases there were blue only, and red only positions which implied that these low energy poses were unique for protein only and glycoprotein model, respectively. Except for Doxorubicin and Sabarubicin, all the moieties docked near black cpk implying that most have some preference for the pocket preferred by ligands in crystal structures. Of course, the extent of the differences varied for each ligand but considering multi-site to single-site interaction hypothesis, the use of glycoprotein model for docking provides new cues.

## Conclusions

Solving reliable and usable structural models of heavily glycosylated proteins remains a challenge in structural biophysics. Human AGP is one such glycoprotein which is of biomedical relevance, key player in drug circulation and its structural information to date is restricted to its protein only form. Deciphering the information by high resolution techniques suffer from genetic basis for isoforms of protein and their varied levels in different racial dispositions and health conditions, and high degree of constitutional and branching polydispersity in the glycans which modify the protein. Sugar moieties differ from proteinogenic portion in solubility criteria, so getting diffraction quality crystals or refinable maps in cryo-EM are still a challenge. The molecular mass of AGP is also too less for solving its map from cryo-EM without complexing it to some Fabs or mAbs. Several attempts have been made to indirectly access the glycan structures by mass spectroscopy, but constructing a usable structure(s) from MS data of AGP has not succeeded to date. Theoretical methods require whetting with experimental structural data which again is limiting. Solution SAXS is a low-resolution technique which can acquire decipherable data from glycoproteins and can provide weighted average information about the molecular shape in solution ^74,75^. If the molecular association remains similar and conformational monodispersity supports globular nature, then chemical polydispersity can be overcome largely in SAXS data analysis.

To test whether this proposition is applicable to human AGP, we acquired SAXS data on purified sample of AGP. Initial SAXS data analysis revealed inherent interparticulate effect at low temperature which eased off upon heating the sample, and monomeric AGP molecules seem to adopt the Dmax and R_g_ close to 6.5-7 and ~2.5 nm, respectively. The protein only portion could be modelled, but expectedly, it was significantly smaller with D_max_ and R_g_ of about 6.3 and 1.7 nm, respectively and its computed SAXS profile did not agree with experimental SAXS datasets. We used previously reported MS profile of glycans and selected the closest possible glycan motif from GLYPROT database to generate four models of fully glycosylated models of AGP. Importantly, theoretical SAXS profiles of all four models of glycoprotein agreed much better to acquired data supporting that these models reliably represent the solution structure of AGP. Realizing that interaction of drug candidates to AGP and albumin are utilized to predict release profiles of the drug, we explored if the glycosylated model could be better or additional template for *in silico* docking /interaction studies? Cavity analysis suggested additional pockets in the glycosylated model of AGP and docking of different drugs showed additional low energy poses composed of protein-glycan constitution of AGP. This suggests that there could be new information which can be computed and experimentally tested to fine tune the protocol of drug release prediction, particularly involving AGP. Overall, we provide a methodology to construct SAXS data based glycosylated models of glycoproteins. As part of continued efforts, we would explore ways to reliably perform molecular dynamics studies with glycosylated version of AGP bound to drug molecules to narrow down differences with experimentally obtained binding data.

## Supporting information

None

## Acknowledgements

We acknowledge 12 FYP project UNSEEN from CSIR which established the SAXS facility in CSIR-IMTECH. Funds from OLP156 were used to procure chemicals. Authors acknowledge faculty and staff for their support. This is IMTECH communication number 29/2023.

## Additional Information

The SAXS datasets for AGP at 283 and 343 K, (and solved 3D models of AGP) are available at SASBDB database under submission IDs SASDPG4 and SASDPH4, respectively.

## Author Contributions

NK – Did final dialysis, VTSAXS experiments, processed data, initial analysis, script writing/review; NP– Initial planning, experiments, attempted to solve information, script review; KP – Tried to place glycanmoieties with earlier data, script review; A – Conceived approaches, analyzed all stages of data, final analysis, script writing.

## Notes

https://www.sasbdb.org/data/SASDPG4/

https://www.sasbdb.org/data/SASDPH4/

### Competing Interest Statement

The authors have declared no competing interest.

