## Supplementary material for "SAXS DATA BASED GLYCOSYLATED MODELS OF HUMAN ALPHA-1-ACID GLYCORPROTEIN, A KEY PLAYER IN HEALTH, DISEASE AND DRUG CIRCULATION": None

**Supplementary Table T1**
Details of SAXS data collection to processing are listed below.

| **Function / Details** | **Program / Details** |
|---|---|
| Instrument/data processing | SAXSpace Programs (Anton Paar) |
| Wavelength of X-rays (nm) | 0.1541 (CuKα) |
| Beam size (μm) | 1 cm*0.4μm (Slit Line Collimation) |
| Camera length (m) | 0.31706 m |
| *Detector* | Mythen 1D (Dectris) |
| *Data Collection Software and Machine Controls* | SAXSDrive Program |
| *Beam Position Correction* | SAXSTreat Program |
| *s* measurement range ($nm^{-1}$) | 0.08–7.5 |
| *Desmearing Software* | SAXSQuant Program using beam profile |
| *Buffer Subtraction* | SAXSQuant Program |
| *Smoothening* | None |
| Monitoring for radiation damage | Flux is too low for radiolysis; Routinely migrationpattern in SDS-PAGE was done |
| Exposure time and frames | Protein: 60 minutes * One frames Matched Buffers: 60 minutes * One frame |
| Sample configuration | Thermostated quartz capillary for line source |
| Sample temperature (K) | 283 – 343 K with increments of 10 K |
| Data Analysis Suite of Programs | ATSAS 3.0.1 and 3.0.2 |
| SASBDB Submissions | SASDPG4 and SASDPH4 |

**Supplementary Figure S1:** Solution SAXS datasets acquired from sample of AGP are presented here. (A) Log-Linear plots of the SAXS datasets at temperatures 283 to 343 K. (B) SAXS datasets of AGP sample as temperature of data collection was decreased from 343 to 283 K. Temperature of experiment are mentioned in the legends. Images of plots were made using ATSAS 3.0.2 Data analysis software.

A

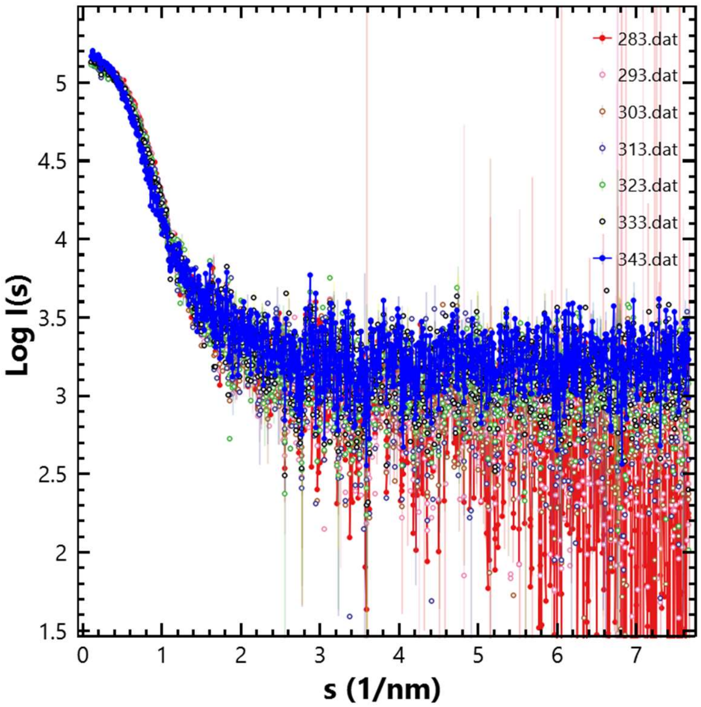


B

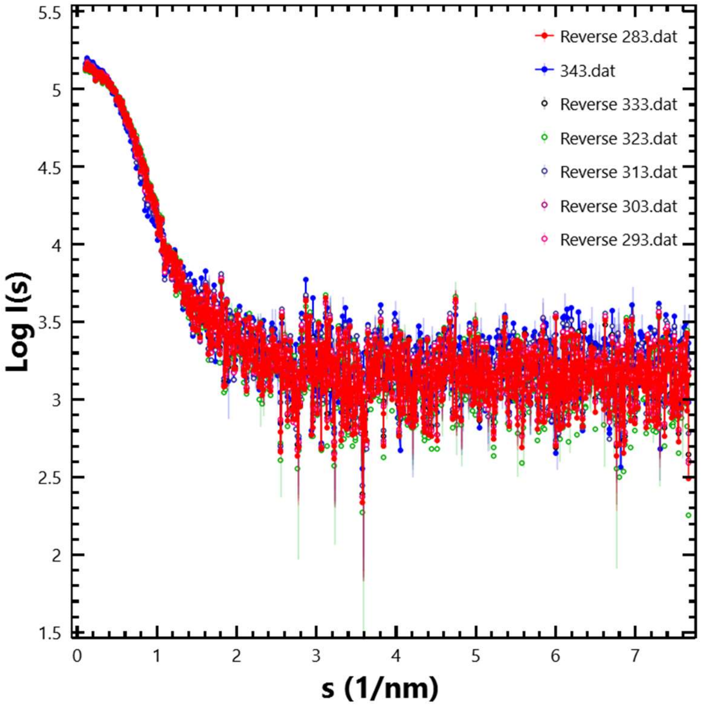

**Supplementary Figure S2:** Guinier and Distance Distribution estimations of the SAXS datasets are presented here as the sample was heated from 283 to 343K. (The experimental temperature of the dataset is mentioned above each set).

For all the datasets, Guinier plot and its corresponding normalized Kratky plot is shown in the top panel. Red line is the Guinier fit and the black line is drawn to guide the eye of reader about the trend of the data beyond the low s region.

The lower panels show the automated estimation of the P(r) profiles. Red line is the fit to the data during the estimation of the interatomic vectors. All images of the analyses are screenshots of the Data Analysis software in ATSAS suite of programs v 3.0.2.

283K [Forward]

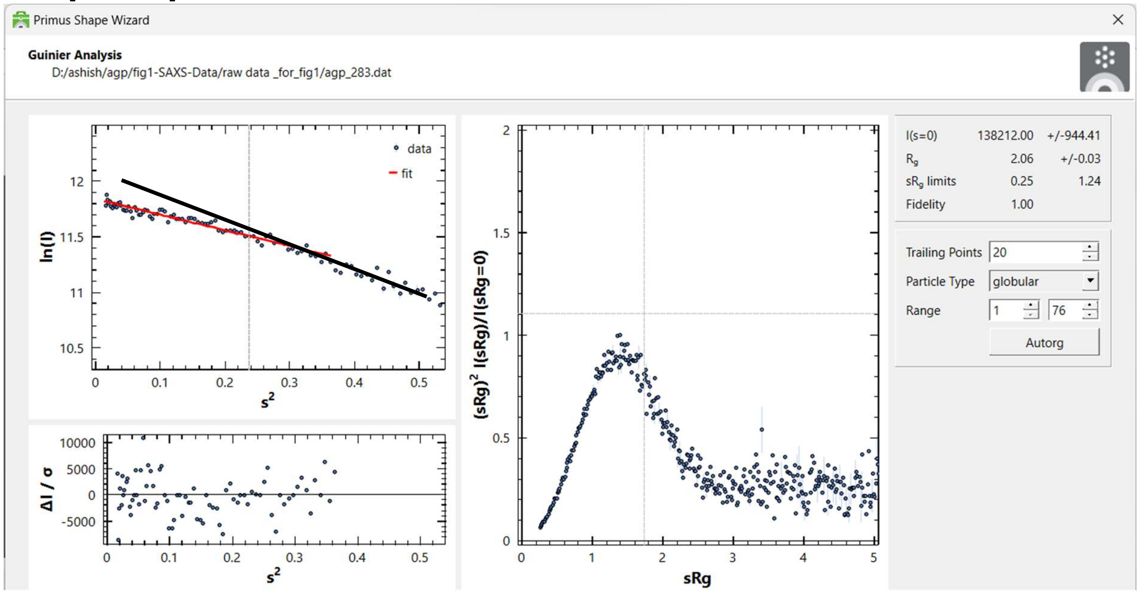


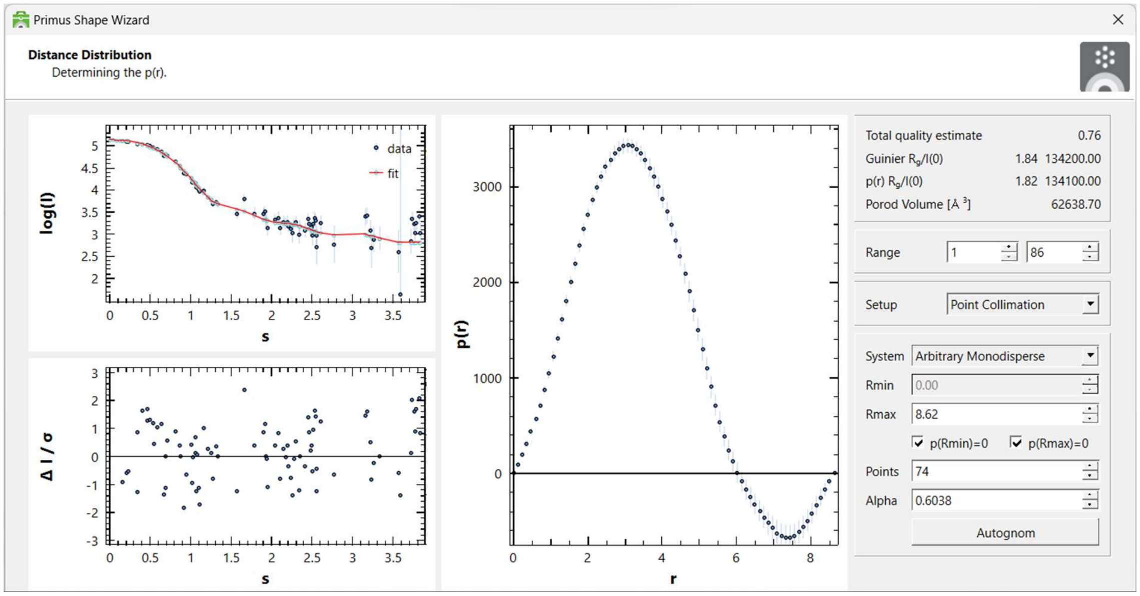

293K [Forward]

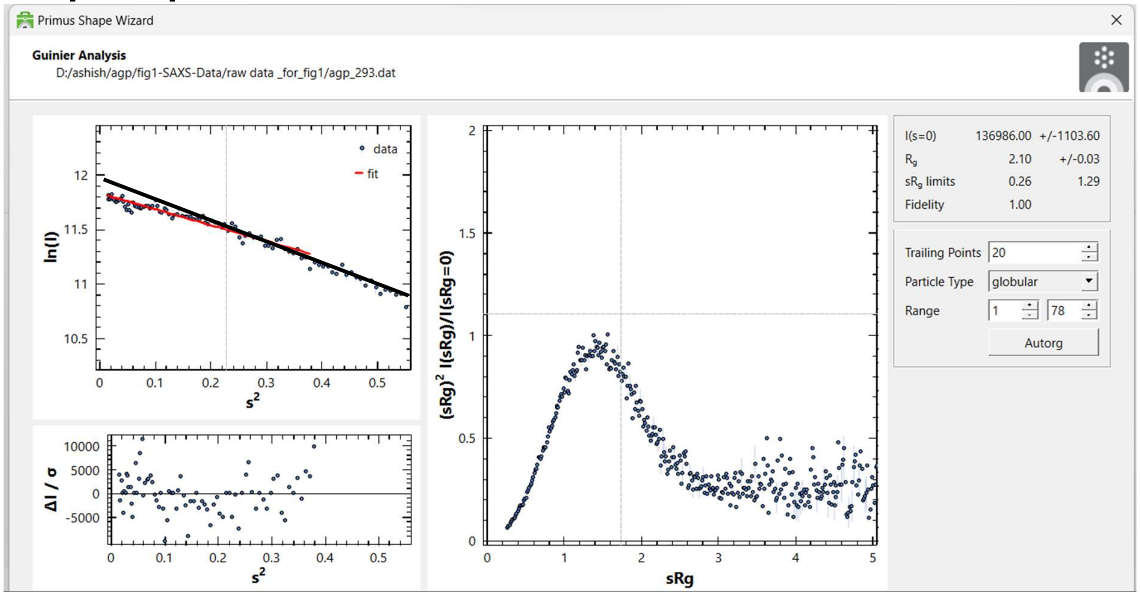


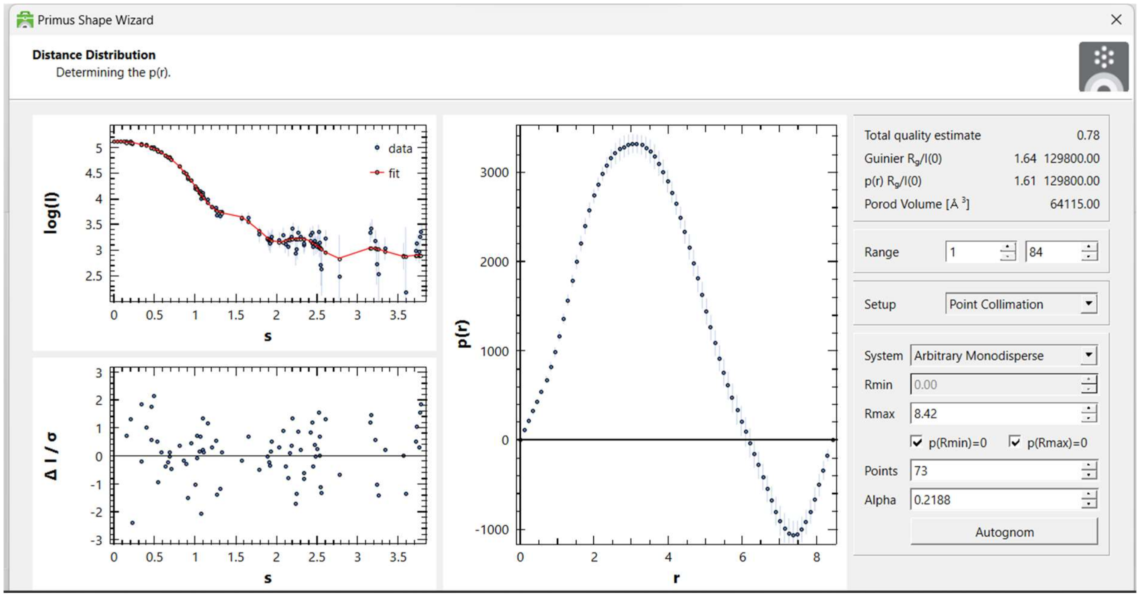

303K [Forward]

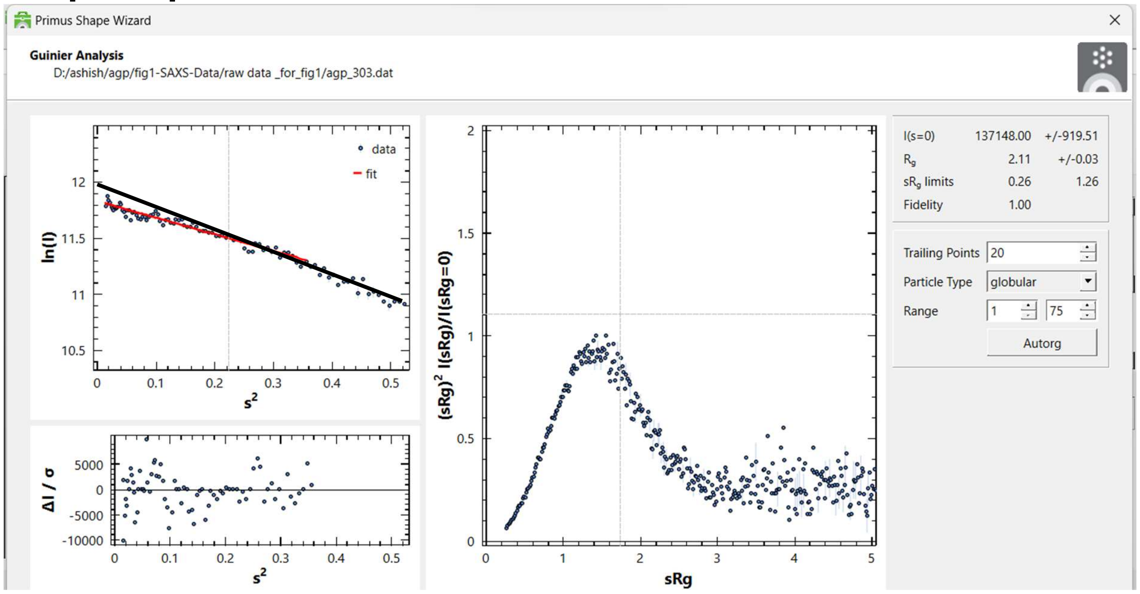


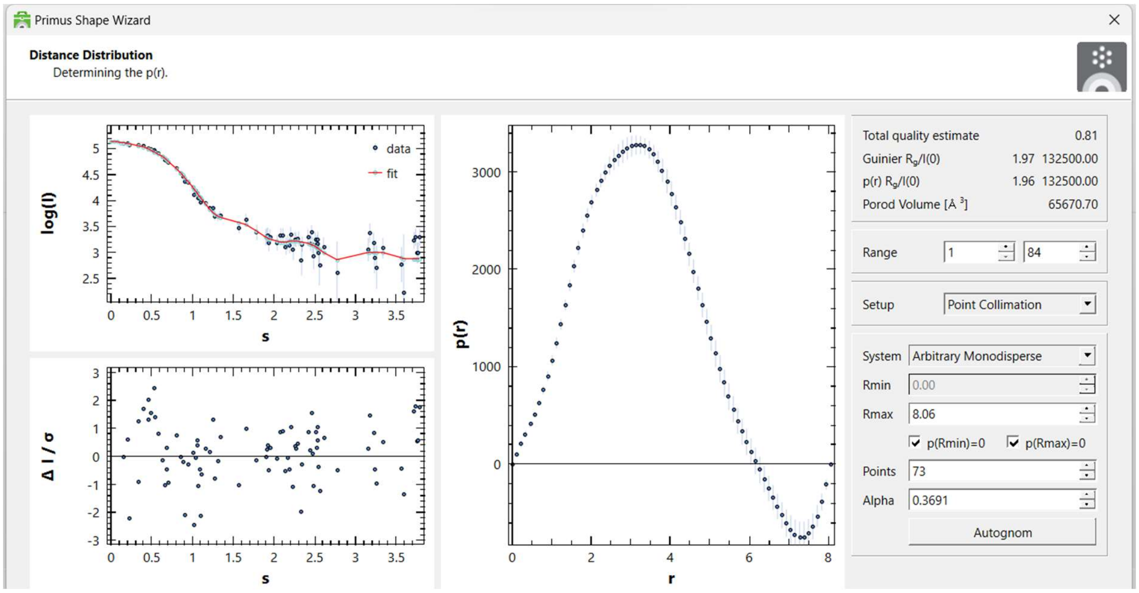

313 K [Forward]

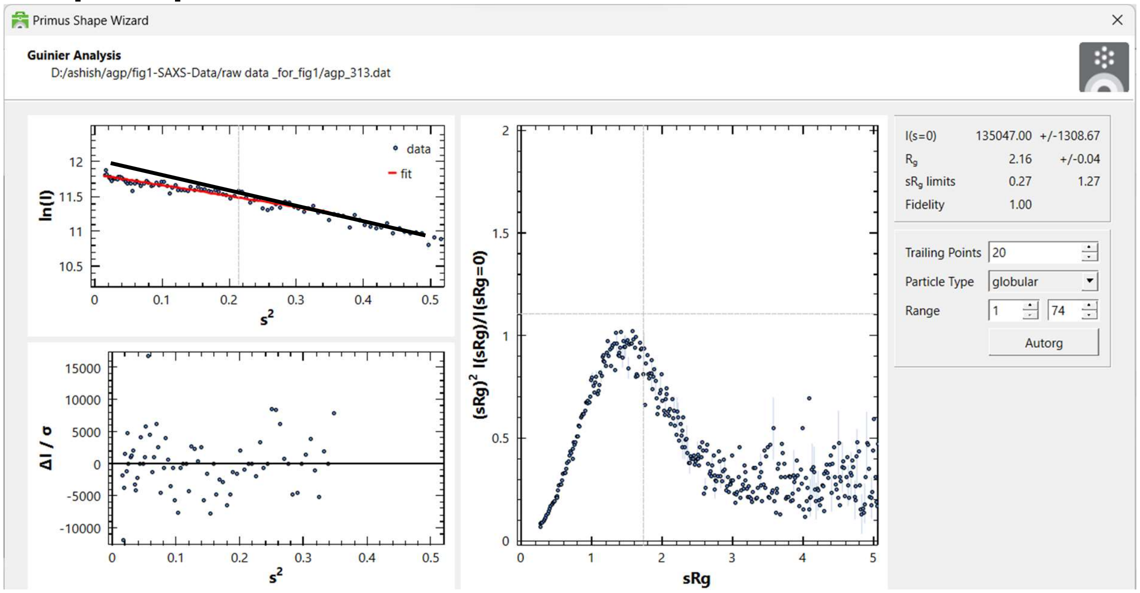


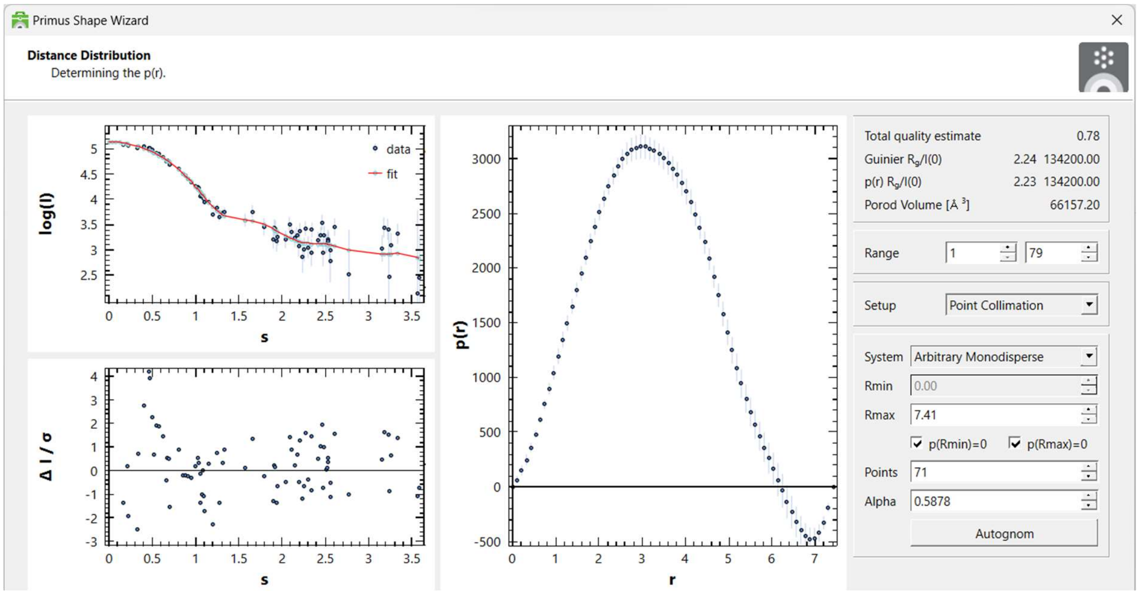

323K [Forward]

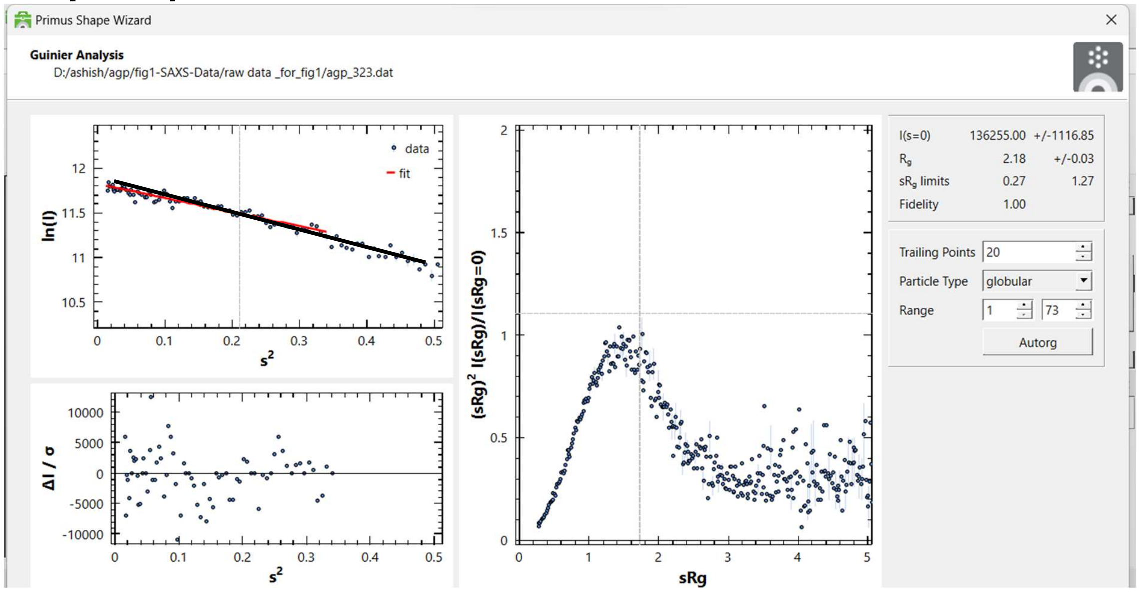


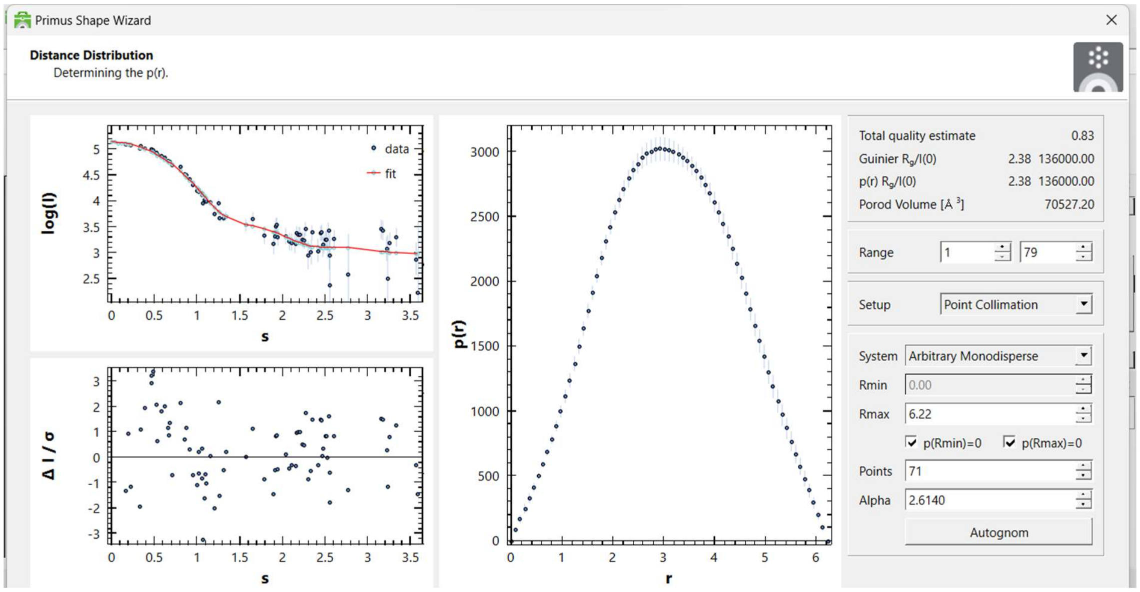

333K [Forward]

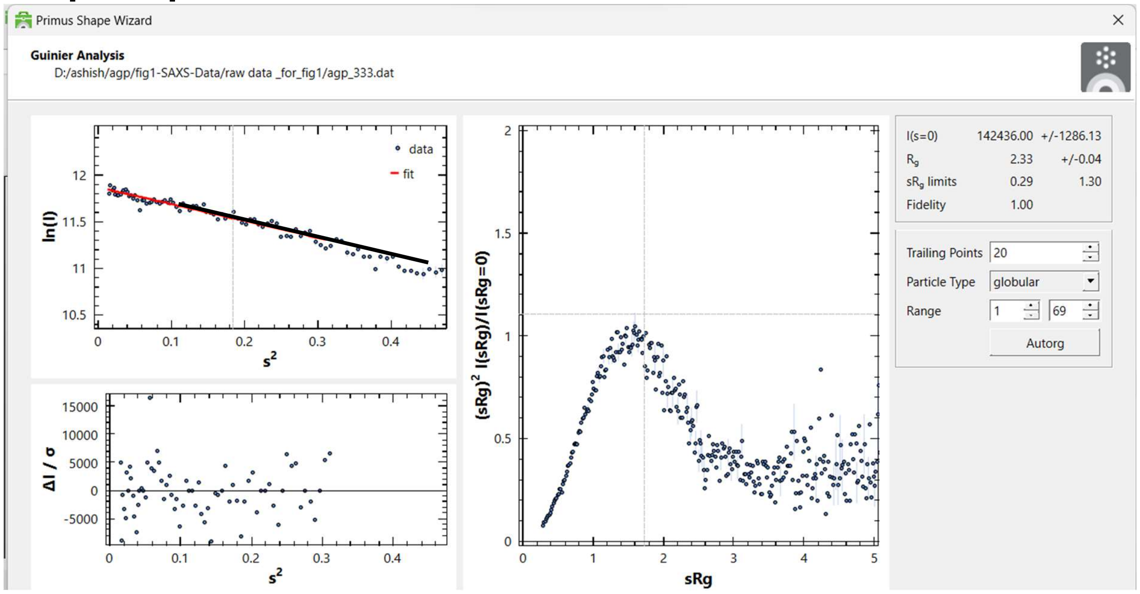


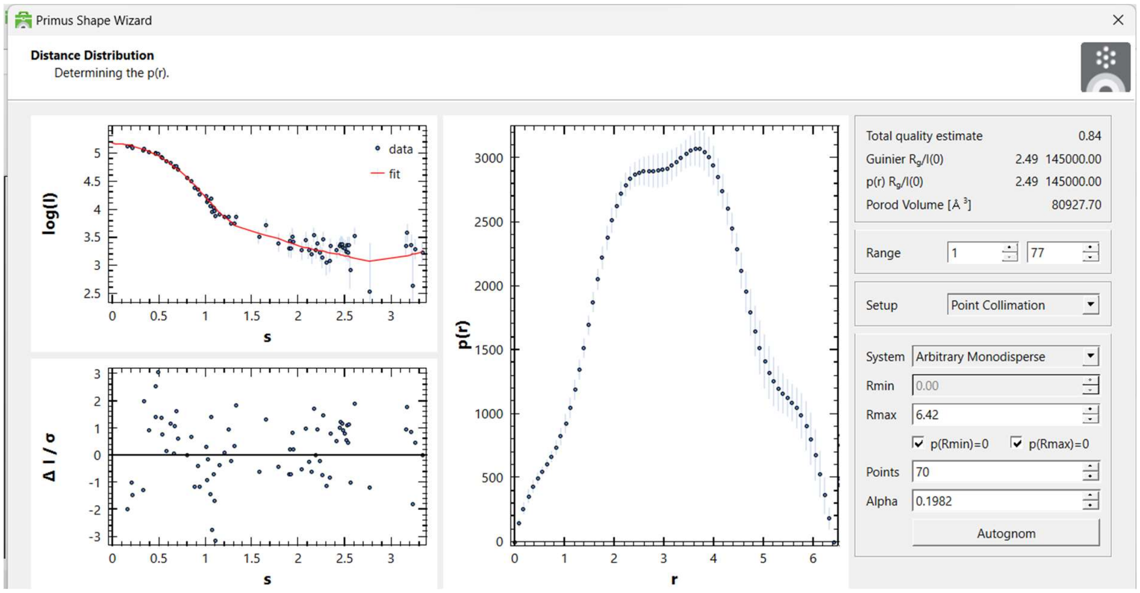

343 K [Forward]

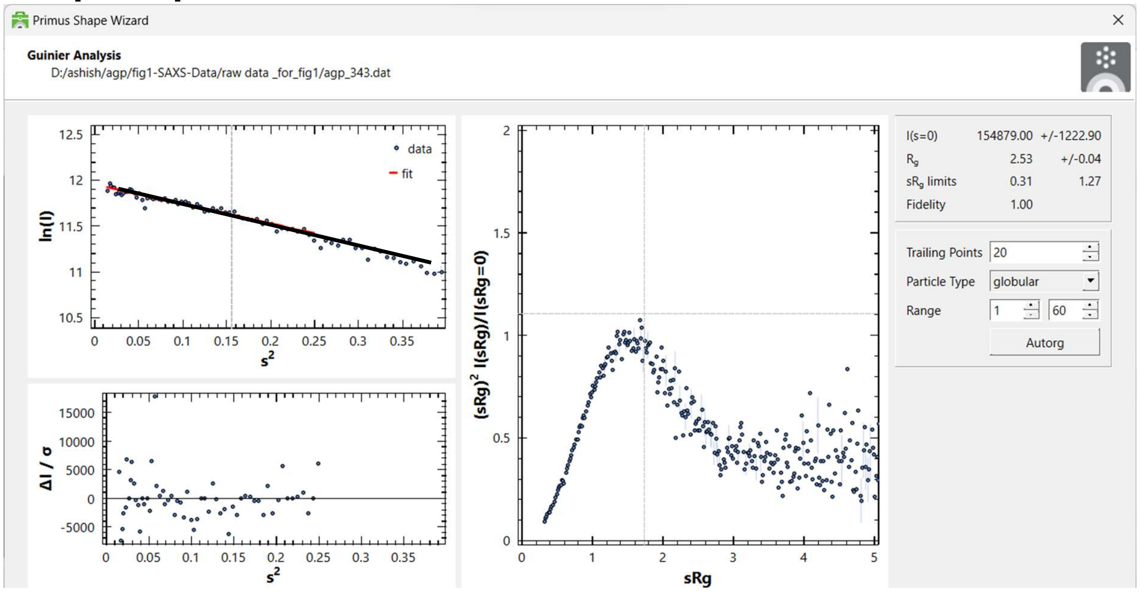


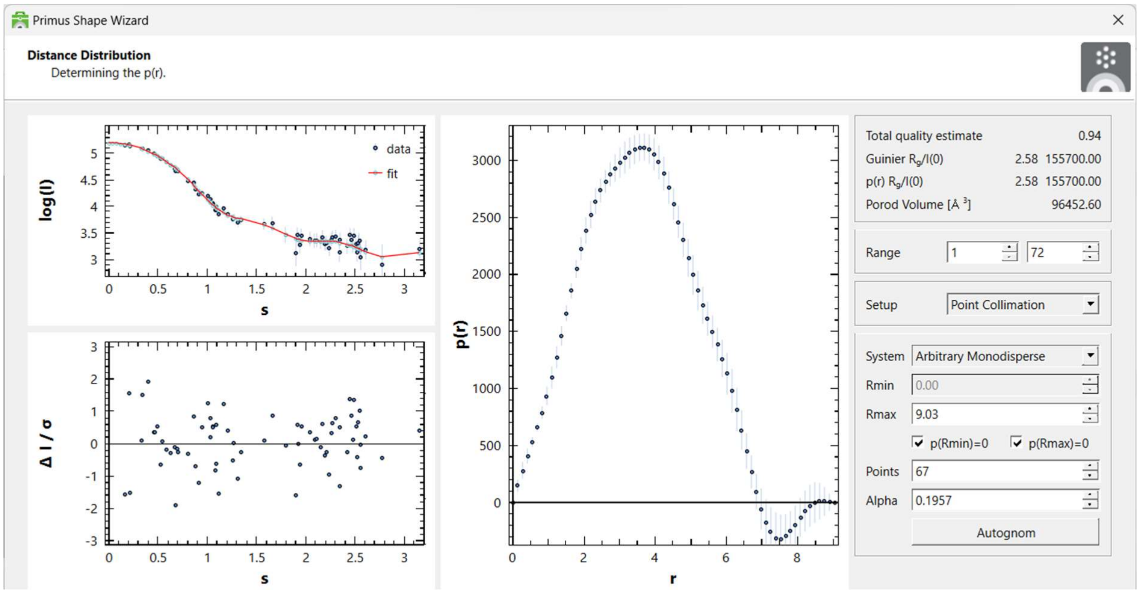

**Supplementary Figure S3:** Crystal structures of AGP solved in unliganded and complexed states. (A) Different crystal structures are shown with their respective PDB ID and ligand mentioned below. The last turns of the C-terminal helix are shown in orange. (B) All five structures are superimposed on each other, and red circle highlights the same pocket The C-terminal helix is shown in orange color.

A

Crystal Structures of A variant of AGP

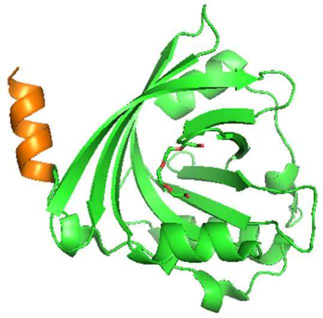

3APU Unliganded A Form

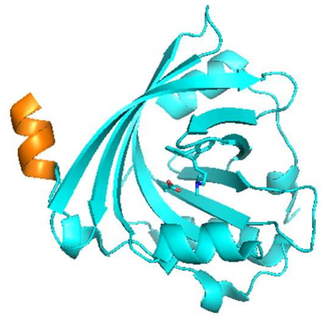

3APV +Amitriptyline

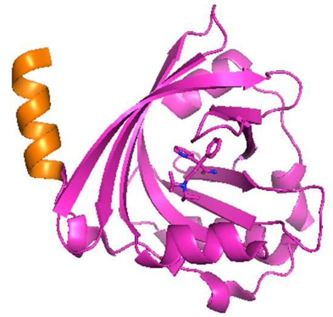

3APW +Disopyramide

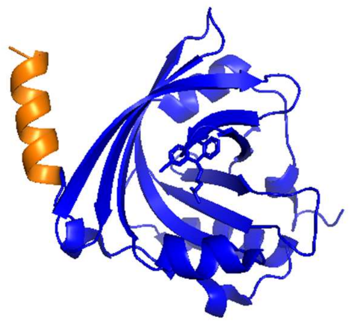

3APX +Chlorpromazine

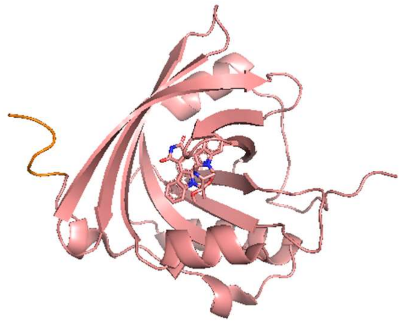

7OUB +UCN01

B

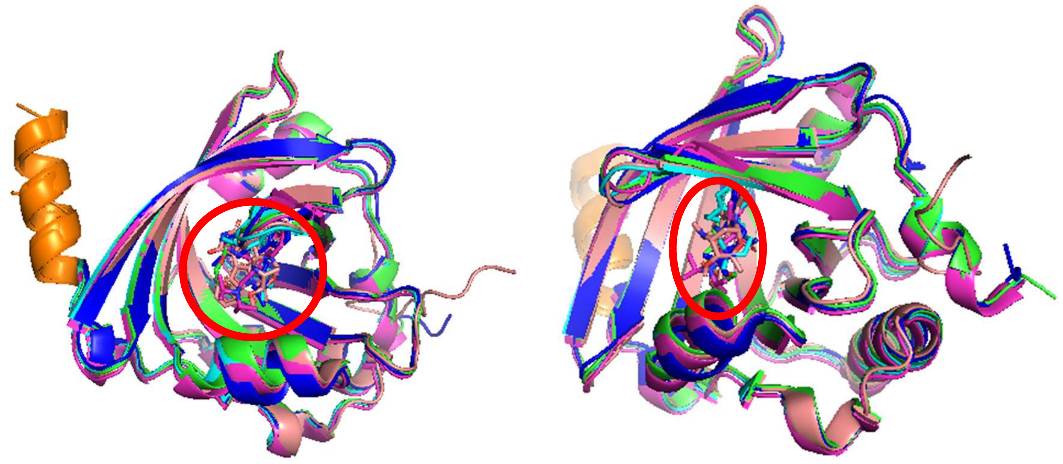

**Supplementary Figure S4**

Results from ALPHAFOLD2 server are shown. (A) and (B) are sequence coverage and predicted Local Distance Difference Test (lDDT) for the template selection. (C) Two rotated views of the five top rankedmodels are shown in cartoon format. The coloring of blue to red was selected for N- to C-terminal representation. The images were generated using PyMoL program. (D) Computed SAXS profiles of thefive models (Lines) predicted by ALPHAFOLD2 server are compared with SAXS datasets at 283 K (Left) and 343 K (Right). Images of plots were made using ATSAS 3.0.1 data analysis software.

A

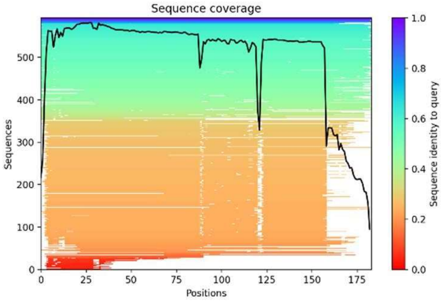


B

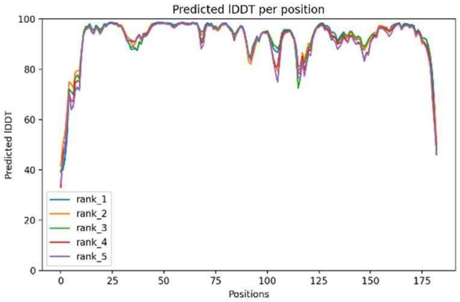


C

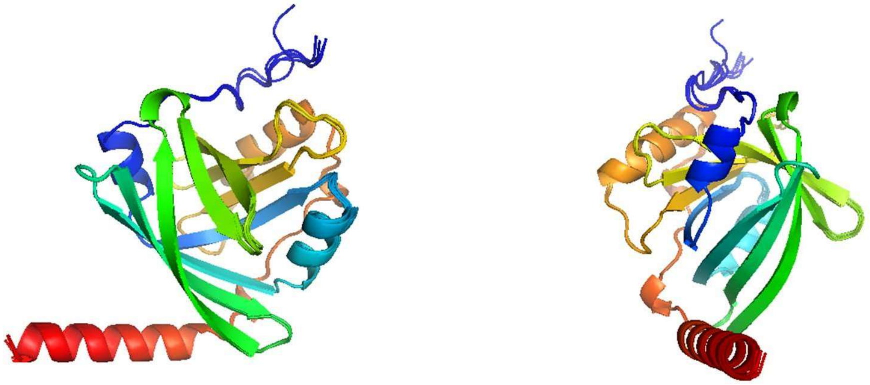

D

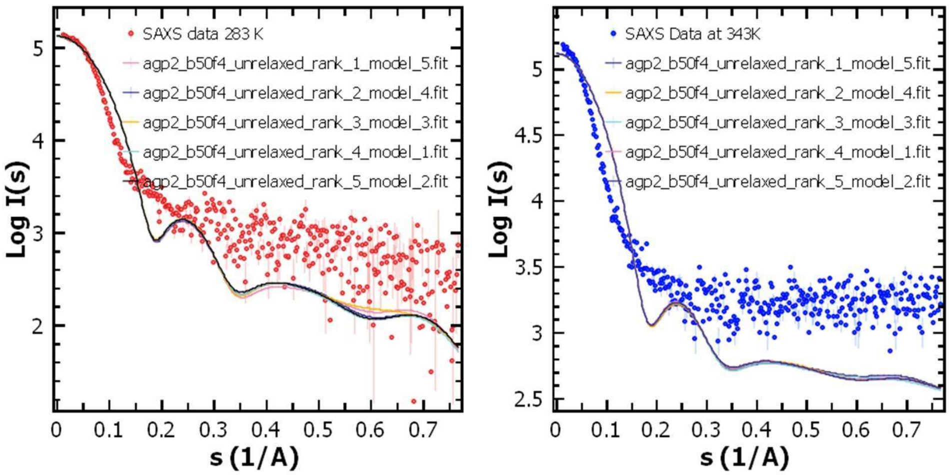

**Supplementary Figure S5**

Glycan motifs selected from GLYPROT database are detailed. Motif number, Position to be connected in protein, their constitution and mass are mentioned. These were guided by results described in Treuheit, M. J., Costello, C. E., & Halsall, H. B. (1992). Analysis of the five glycosylation sites of human alpha 1-acid glycoprotein. Biochem J, 283 ( Pt 1), 105-112. https://doi.org/10.1042/bj2830105.

GLYPROT server and details: http://www.glycosciences.de/modeling/glyprot/php/main.php

*__Position 1__*

Glycan Motif 8959

```
a-D-Neup5Ac-(2-3)-b-D-Galp-(1-4)-b-D-GlcpNAc-(1-3)-b-D-Galp-(1-4)-b-D-GlcpNAc-(1-6)+
                                                                                   |
                                                                              a-D-Manp-(1-6)+            a-L-Fucp-(1-6)+
                                                                                   |        |                          |
                                 a-D-Neup5Ac-(2-3)-b-D-Galp-(1-4)-b-D-GlcpNAc-(1-2)+        |                  b-D-GlcpNAc
                                                                                            |                          |
                                                                                       b-D-Manp-(1-4)-b-D-GlcpNAc-(1-4)+
                                                                                            |
                                 a-D-Neup5Ac-(2-3)-b-D-Galp-(1-4)-b-D-GlcpNAc-(1-4)+        |
                                                                                   |        |
                                                                              a-D-Manp-(1-3)+
                                                                                   |
                                 a-D-Neup5Ac-(2-3)-b-D-Galp-(1-4)-b-D-GlcpNAc-(1-2)+
```

4029 Da

Glycan Motif 8533

```
a-D-Neup5Ac-(2-3)-b-D-Galp-(1-4)-b-D-GlcpNAc-(1-3)-b-D-Galp-(1-4)-b-D-GlcpNAc-(1-6)+
                                                                                   |
                                                                              a-D-Manp-(1-6)+            a-L-Fucp-(1-6)+
                                                                                   |        |                          |
                                 a-D-Neup5Ac-(2-3)-b-D-Galp-(1-4)-b-D-GlcpNAc-(1-2)+        |                  b-D-GlcpNAc
                                                                                            |                          |
                                                                                       b-D-Manp-(1-4)-b-D-GlcpNAc-(1-4)+
                                                                                            |
                                 a-D-Neup5Ac-(2-3)-b-D-Galp-(1-4)-b-D-GlcpNAc-(1-4)+        |
                                                                                   |        |
                                                                              a-D-Manp-(1-3)+
                                                                                   |
                                 a-D-Neup5Ac-(2-3)-b-D-Galp-(1-4)-b-D-GlcpNAc-(1-2)+
```

4029 Da

*__Position 2__*

Glycan Motif 9119

```
a-D-Neup5Ac-(2-6)-b-D-Galp-(1-4)-b-D-GlcpNAc-(1-2)-a-D-Manp-(1-6)+
                                                                 |
         a-D-Neup5Ac-(2-3)-b-D-Galp-(1-4)-b-D-GlcpNAc-(1-4)-b-D-Manp-(1-4)-b-D-GlcpNAc-(1-4)-b-D-GlcpNAc
                                                                 |
a-D-Neup5Ac-(2-6)-b-D-Galp-(1-4)-b-D-GlcpNAc-(1-2)-a-D-Manp-(1-3)+
```

2862 Da

Glycan Motif 9085

```
a-D-Neup5Ac-(2-3)-b-D-Galp-(1-4)-b-D-GlcpNAc-(1-2)-a-D-Manp-(1-6)+
                                                                 |
      a-D-Neup5Ac-(2-6)-b-D-Galp-(1-4)-b-D-GlcpNAc-(1-6)+   b-D-Manp-(1-4)-b-D-GlcpNAc-(1-4)-b-D-GlcpNAc
                                                        |        |
                                                   a-D-Manp-(1-3)+
                                                        |
      a-D-Neup5Ac-(2-6)-b-D-Galp-(1-4)-b-D-GlcpNAc-(1-2)+
```

2862 Da

Glycan Motif 9073

```
      a-D-Neup5Ac-(2-6)-b-D-Galp-(1-4)-b-D-GlcpNAc-(1-6)+
                                                        |
                                                   a-D-Manp-(1-6)+
                                                        |        |
      a-D-Neup5Ac-(2-3)-b-D-Galp-(1-4)-b-D-GlcpNAc-(1-2)+   b-D-Manp-(1-4)-b-D-GlcpNAc-(1-4)-b-D-GlcpNAc
                                                                 |
a-D-Neup5Ac-(2-6)-b-D-Galp-(1-4)-b-D-GlcpNAc-(1-2)-a-D-Manp-(1-3)+
```

2862 Da

*Position 3*

Glycan Motif 8951

```
a-D-Neup5Ac-(2-3)-b-D-Galp-(1-4)-b-D-GlcpNAc-(1-3)-b-D-Galp-(1-4)-b-D-GlcpNAc-(1-6)+
                                                                                   |
                                                                              a-D-Manp-(1-6)+            a-L-Fucp-(1-6)+
                                                                                   |        |                          |
a-D-Neup5Ac-(2-3)-b-D-Galp-(1-4)-b-D-GlcpNAc-(1-3)-b-D-Galp-(1-4)-b-D-GlcpNAc-(1-2)+        |                  b-D-GlcpNAc
                                                                                            |                          |
                                                                                       b-D-Manp-(1-4)-b-D-GlcpNAc-(1-4)+
                                                                                            |
                                 a-D-Neup5Ac-(2-3)-b-D-Galp-(1-4)-b-D-GlcpNAc-(1-4)+        |
                                                                                   |        |
                                                                              a-D-Manp-(1-3)+
                                                                                   |
                                                   b-D-Galp-(1-4)-b-D-GlcpNAc-(1-2)+
```

4108 Da

*Position 4*

Glycan Motif 9232

```
      a-L-Fucp-(1-3)-b-D-GalpNAc-(1-3)-a-D-Galp-(1-3)-b-D-Galp-(1-4)-b-D-GlcpNAc-(1-6)+
                                                                                      |
a-L-Fucp-(1-3)-b-D-GalpNAc-(1-3)-a-D-Galp-(1-3)-b-D-Galp-(1-4)-b-D-GlcpNAc-(1-4)-a-D-Manp-(1-6)+
                                                                                      |        |
      a-L-Fucp-(1-3)-b-D-GalpNAc-(1-3)-a-D-Galp-(1-3)-b-D-Galp-(1-4)-b-D-GlcpNAc-(1-2)+        |
                                                                                               |
                                                                                          b-D-Manp-(1-4)-b-D-GlcpNAc-(1-4)-b-D-GlcpNAc
                                                                                               |
      a-L-Fucp-(1-3)-b-D-GalpNAc-(1-3)-a-D-Galp-(1-3)-b-D-Galp-(1-4)-b-D-GlcpNAc-(1-4)+        |
                                                                                      |        |
                                                                                 a-D-Manp-(1-3)+
                                                                                      |
      a-L-Fucp-(1-3)-b-D-GalpNAc-(1-3)-a-D-Galp-(1-3)-b-D-Galp-(1-4)-b-D-GlcpNAc-(1-2)+
```

5274 Da

Position 5

Glycan Motif 8996

```
a-D-Neup5Ac-(2-3)-b-D-Galp-(1-4)-b-D-GlcpNAc-(1-6)+
                                                  |
                                             a-D-Manp-(1-6)+            a-L-Fucp-(1-6)+
                                                  |        |                          |
a-D-Neup5Ac-(2-6)-b-D-Galp-(1-4)-b-D-GlcpNAc-(1-2)+        |                  b-D-GlcpNAc
                                                           |                          |
                                                      b-D-Manp-(1-4)-b-D-GlcpNAc-(1-4)+
                                                           |
a-D-Neup5Ac-(2-3)-b-D-Galp-(1-4)-b-D-GlcpNAc-(1-4)+        |
                                                  |        |
                                             a-D-Manp-(1-3)+
                                                  |
a-D-Neup5Ac-(2-6)-b-D-Galp-(1-4)-b-D-GlcpNAc-(1-2)+
```

3664 Da

**Supplementary Figure S6**

Results of FPOCKET Web Server [https://bioserv.rpbs.univ-paris-diderot.fr/services/fpocket/] whichshows sequence location of pockets in two models of AGP.

Unglycosylated Model using probe of 6 Å radius

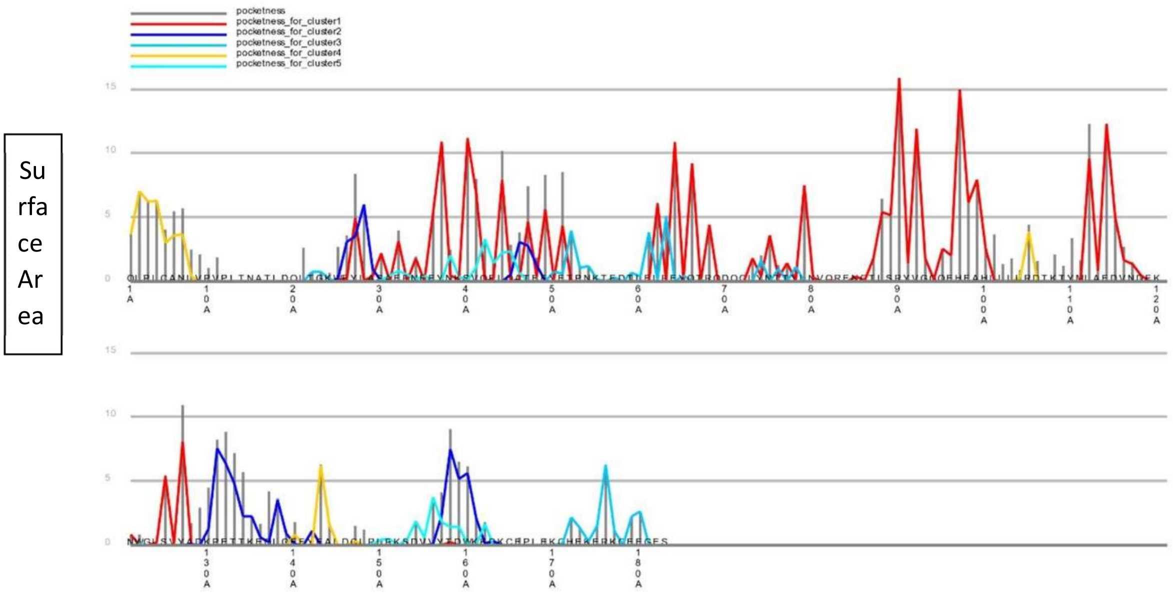


Pockets in Glycosylated Model 1 using probe of 6 Å radius

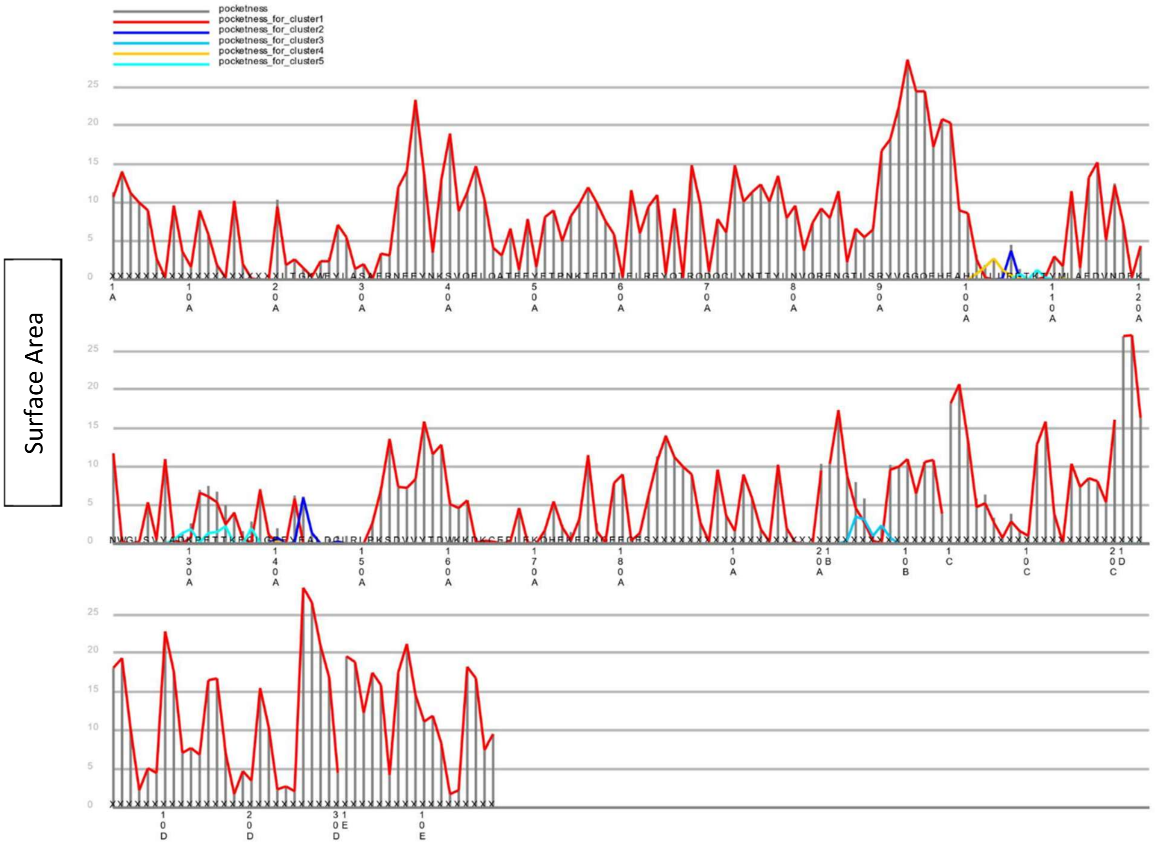

**Supplementary Figure S7**

Pockets identified in the unglycosylated (blue for 6Å and cyan for 10Å) and glycosylated (red for 6Å and salmon for 10Å) model of AGP using GHECOM server. The images were generated using PyMoLprogram.

Using probe of 6 Å radius

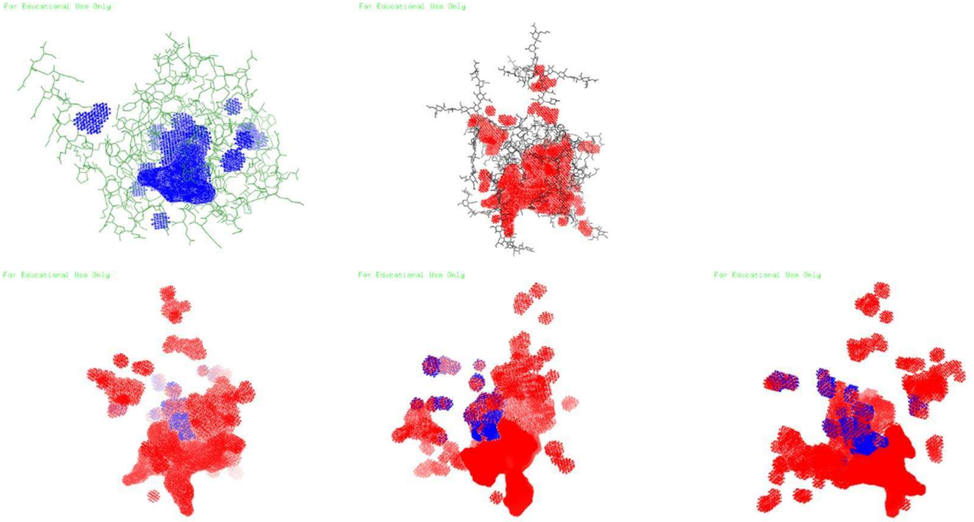

Using probe of 10 Å radius

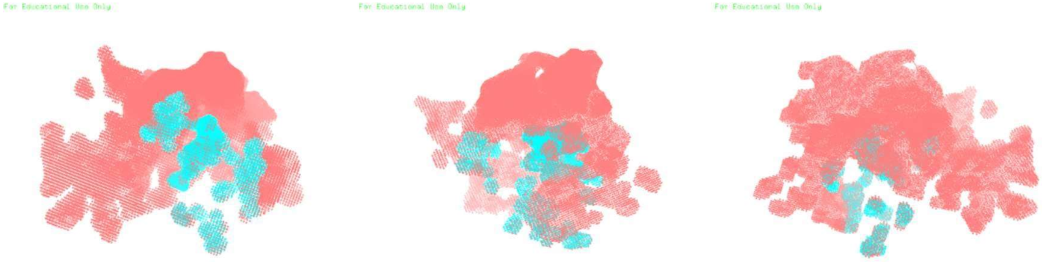

**Supplementary Figure S8:** Two rotated views of the glycoprotein model (grey sticks and backbone in cartoon format) with Disopyramide (black cpk) in same 3D position as in crystal structure (PDB ID: 3APW) are shown here. The C-terminal helix is shown in orange color.

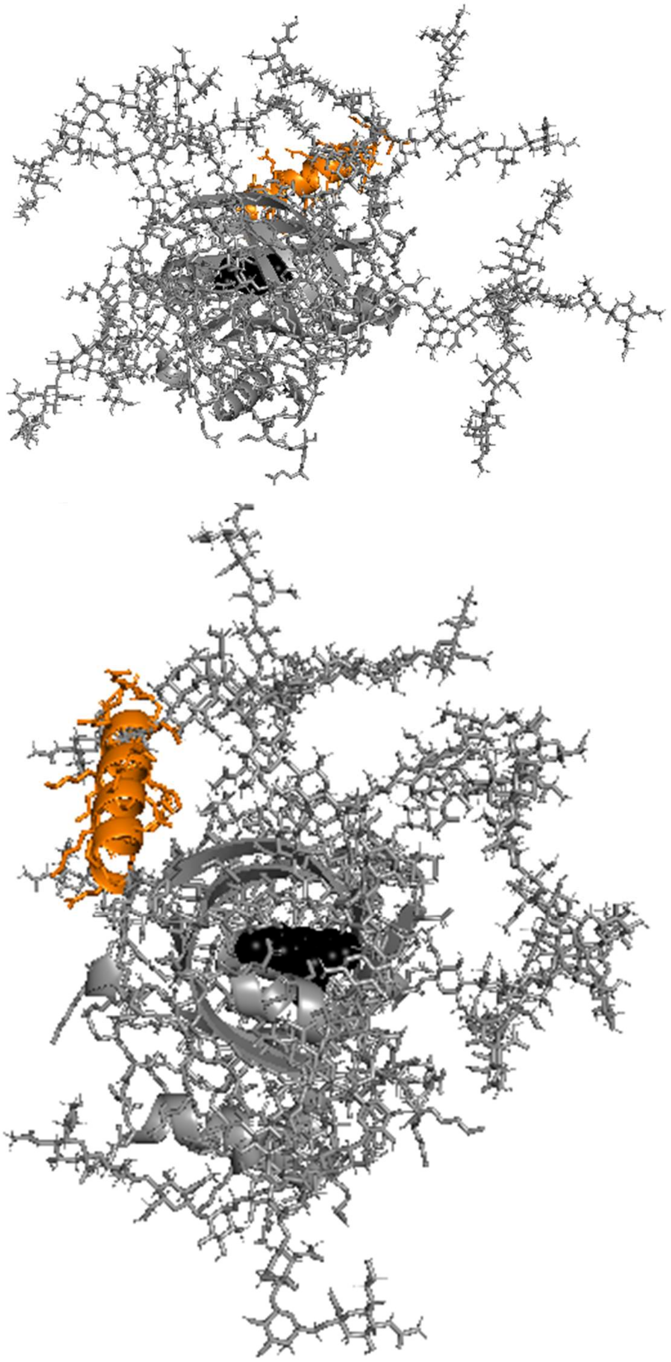

**Supplementary Figure S9:** As support to the images in Figure 5, two rotated views of the glycoprotein model showing top ten poses of the drug molecule (blue docked on protein only model, and red docked on the glycoprotein model). The Disopyramide as seen in crystal structure (PDB ID 3APW). All organic moieties are shown as cpk. As before, the C-terminal helix is shown as orange cartoon. All images have been generated using the PyMol program.

Aripiprazole

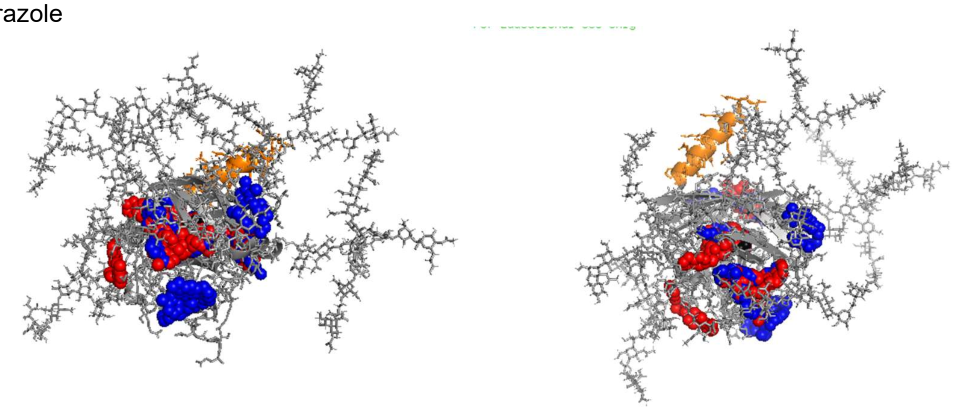

Staurosporine

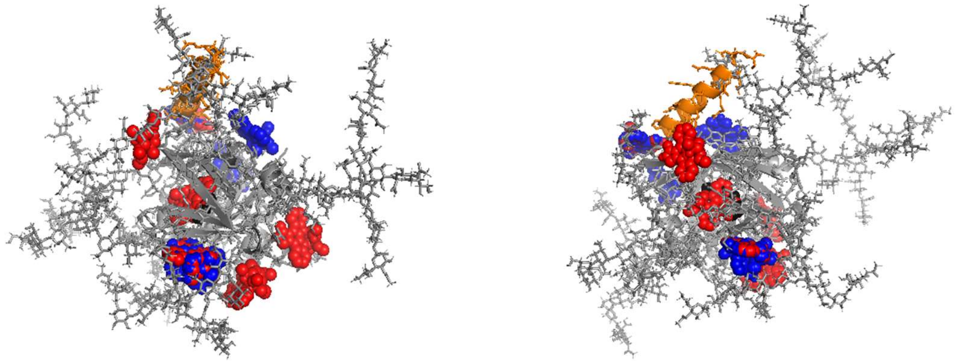

Progesterone

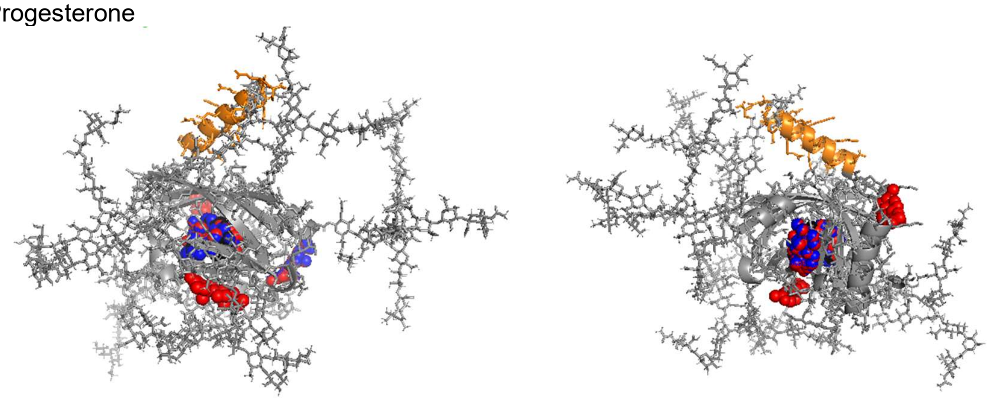

Propranolol

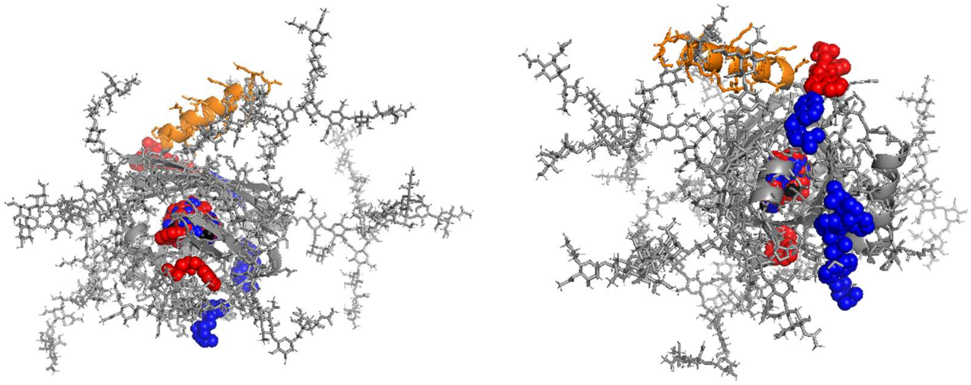

Verapamil

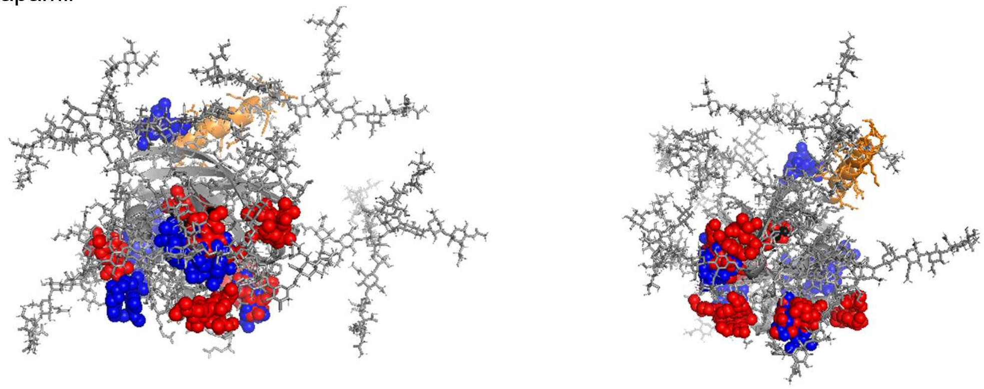

Disopyramide

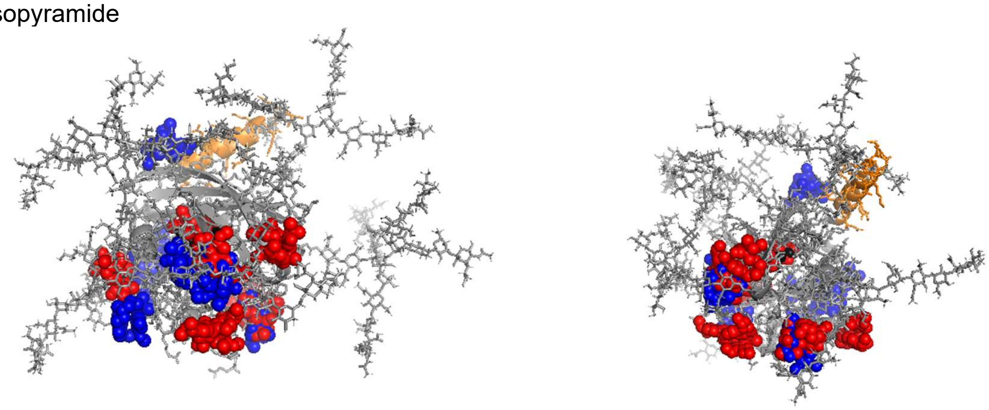

Warfarin

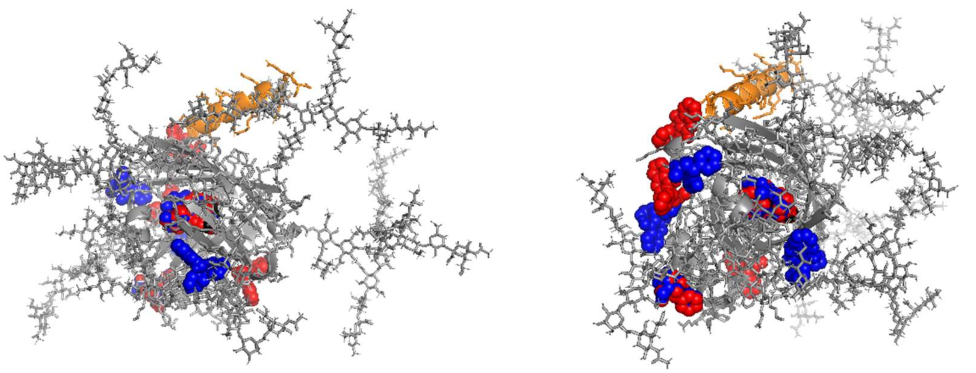

Tuberine

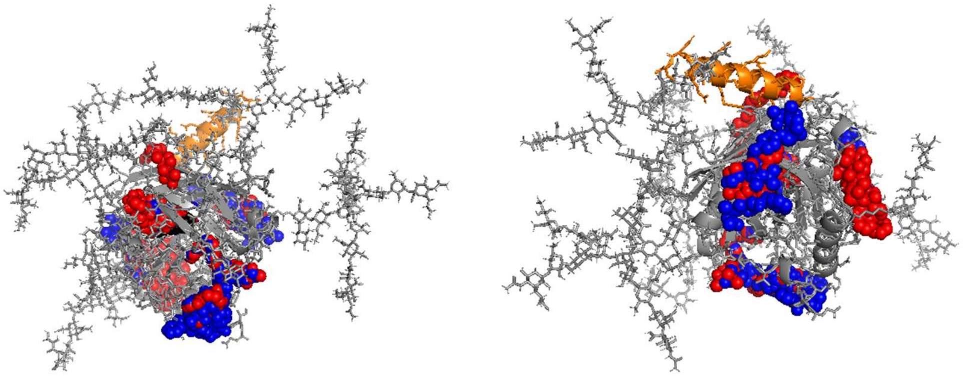

Ostreogrycin A

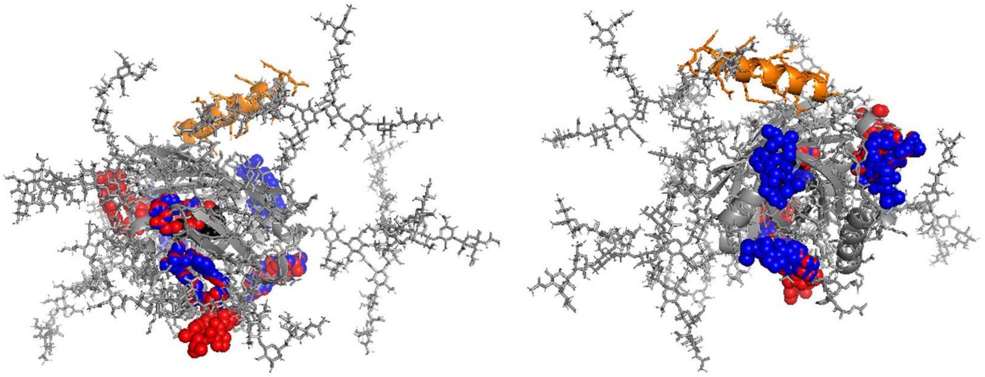

Doxorubicin

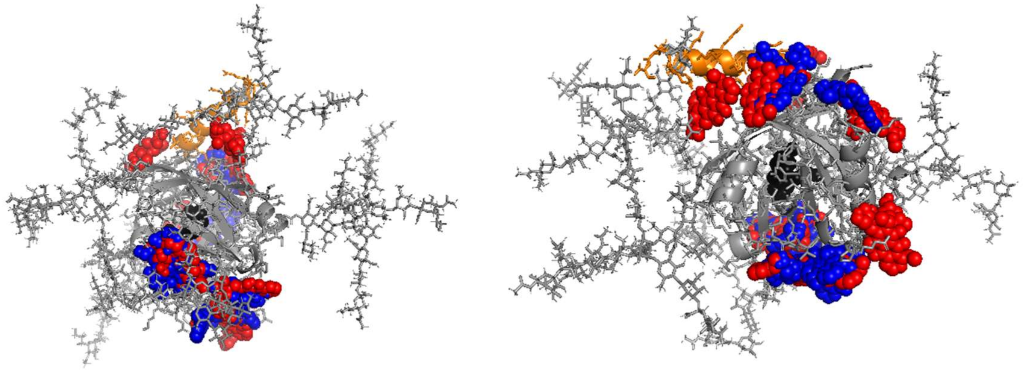

Sabarubicin

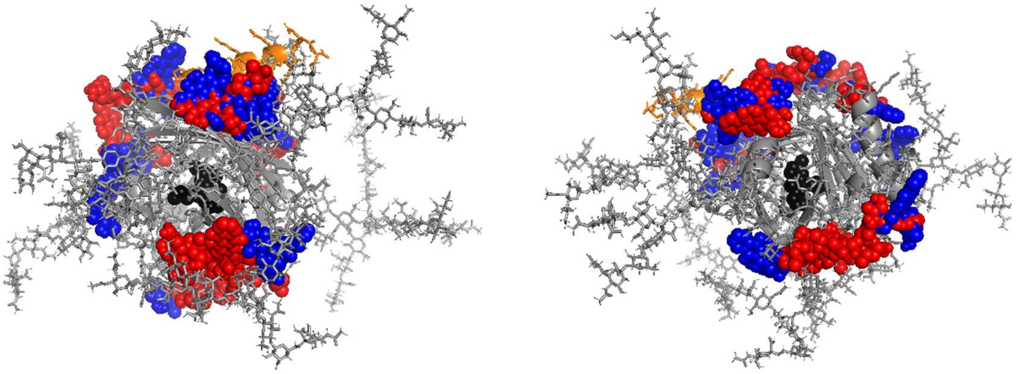

Sucrose

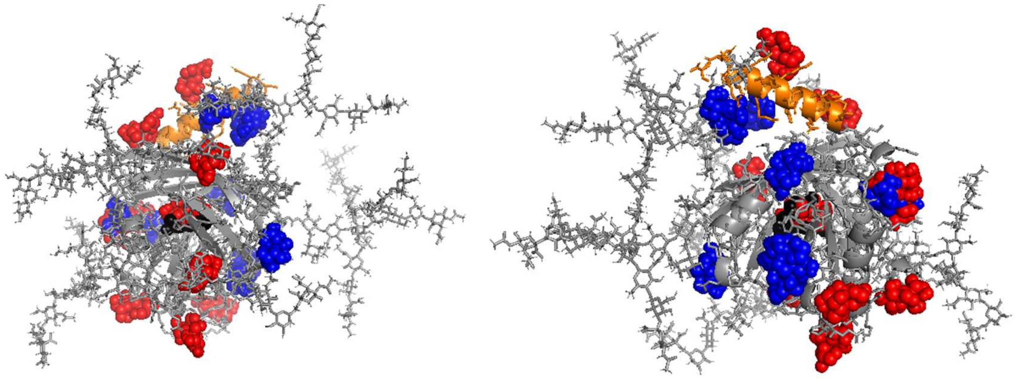
